# ER-associated biogenesis of PINK1 preprotein for neuronal mitophagy

**DOI:** 10.1101/2024.06.21.600039

**Authors:** J. Tabitha Hees, Inmaculada Segura, Andrea Schneider, Martina Schifferer, Thomas Misgeld, Angelika B. Harbauer

## Abstract

A central role in mitochondrial quality control is played by the Parkinson-related mitochondrial kinase PINK1, whose mRNA is transported in neurons by mitochondrial hitch- hiking. Using a live-cell imaging assay for the translation of the PINK1 precursor, we show that local translation of PINK1 requires a concerted interplay between mitochondria and the ER in neurons. For efficient translation, the *Pink1* mRNA needs to relocate to ribosomes located near endolysosomes and the ER. The ER membrane-tethered chaperone DNAJB6 then shields the PINK1 precursor on transit to mitochondria following the ER-SURF pathway. Loss of DNAJB6 hence leads to persistence of ER/endolysosome-associated PINK1 precursor stores and failure of mitophagy upon mitochondrial damage.

## Main text

Mitochondria are of particular importance in neurons, whose vast energetic demand is almost exclusively met by ATP generated through oxidative phosphorylation in mitochondria (*1*). The PTEN-induced kinase 1 (PINK1) is a serine/threonine kinase involved in sensing mitochondrial damage and it is crucial for neuronal health, as point mutations in *PINK1* gene lead to hereditary forms of Parkinson’s Disease (PD) (*2*). Like most nuclear-encoded mitochondrial precursors, PINK1 contains an N-terminal mitochondrial targeting sequence (MTS), which in PINK1 is followed by a transmembrane (TM) domain that is cleaved by the Presenilin-Associated Rhomboid-Like Protein (PARL) (Fig. S1A), allowing the re-export and degradation of cleaved PINK1 when mitochondria are healthy (*3*). In response to mitochondrial damage, however, the PINK1 protein stabilizes at the outer membrane and phosphorylates itself, several outer membrane proteins and ubiquitin (*4–7*), which triggers the translocation of the cytosolic E3-ubiquitin ligase Parkin to the damaged organelle (*8*). This initiates a cascade of ubiquitination and phosphorylation, leading to the demarcation of damaged mitochondria with phospho-ubiquitin chains, recruitment of autophagy adaptors and selective autophagy of the damaged organelle (*5–7*, *9–12*).

Given the intricate mechanism of PINK1 protein import and stabilization (*3*), we reasoned that the site of protein translation may have direct consequences on the ability of mitochondria to process and stabilize PINK1 in response to damage. Transport of *Pink1* mRNA on the mitochondrial surface keeps a steady supply of freshly synthesized PINK1 precursor available in the axon as well as in dendrites (*13*). This attachment is mediated by an RNA anchoring complex on the mitochondrial surface formed by the neuronal RNA binding protein (RBP) Synaptojanin 2a (SYNJ2a) and its binding protein (SYNJ2BP) at the mitochondrial outer membrane. The association of *Pink1* mRNA with mitochondria is repressed by insulin signalling but enhanced by AMP-activated protein kinase (AMPK) signalling, the kinase which directly phosphorylates SYNJ2BP (*14*). Hence, *Pink1* mRNA tethering restricts neuronal mitophagy to periods of high insulin, whose downstream signalling cascade represses AMPK (*14*, *15*). However, untethering of the *Pink1* mRNA from mitochondria, which favors neuronal mitophagy, places a higher distance between the mRNA and the final destination of its translation product with yet to be determined consequences for the cytosolic journey of the PINK1 precursor.

## Visualization and regulation of PINK1 translation

To trace the spatial distribution of PINK1 translation in living neurons, we designed a live-cell imaging reporter for PINK1 based on the SunTag-visualization system (*16*). Briefly, inclusion of a repetitive array of GCN4 epitopes in the PINK1 open reading frame leads to the clustering of co-expressed GFP-tagged nanobodies (scFv-GFP) as soon as the PINK1 nascent chain emerges from the ribosome (Fig. 1A). We compared the placement of the array before the transmembrane segment of PINK1 (PINK1-SunTagmito) with the placement after the transmembrane domain (PINK1-SunTagcyto) (Fig. 1B, S1A). PINK1-SunTagcyto formed small puncta close to mitochondria, which faithfully recapitulated PINK1 biology, as they disappeared quickly after translational inhibition (Fig. 1C-D, S1B). Furthermore, the encoded protein was stabilized by mitochondrial depolarization or proteasome inhibition at the expected molecular weights (Fig. S1C). This was unlike PINK1-SunTagmito, which showed signs of delayed import (sequential cleavage by MPP and PARL (*17*), Fig. S1D) and its signal was still observed upon translation inhibition (Puromycin, Fig. 1C-D, S1B). This was not due to interference of the GFP-tagged nanobody used for imaging, as the delayed cleavage was also evident in the absence of the nanobody (Fig. S1E-F). We therefore selected the PINK1-SunTagcyto reporter (from now on called PINK1-SunTag) to test the effect of untethering of the *Pink1* mRNA by inhibition of AMPK signaling.

**Fig. 1.**
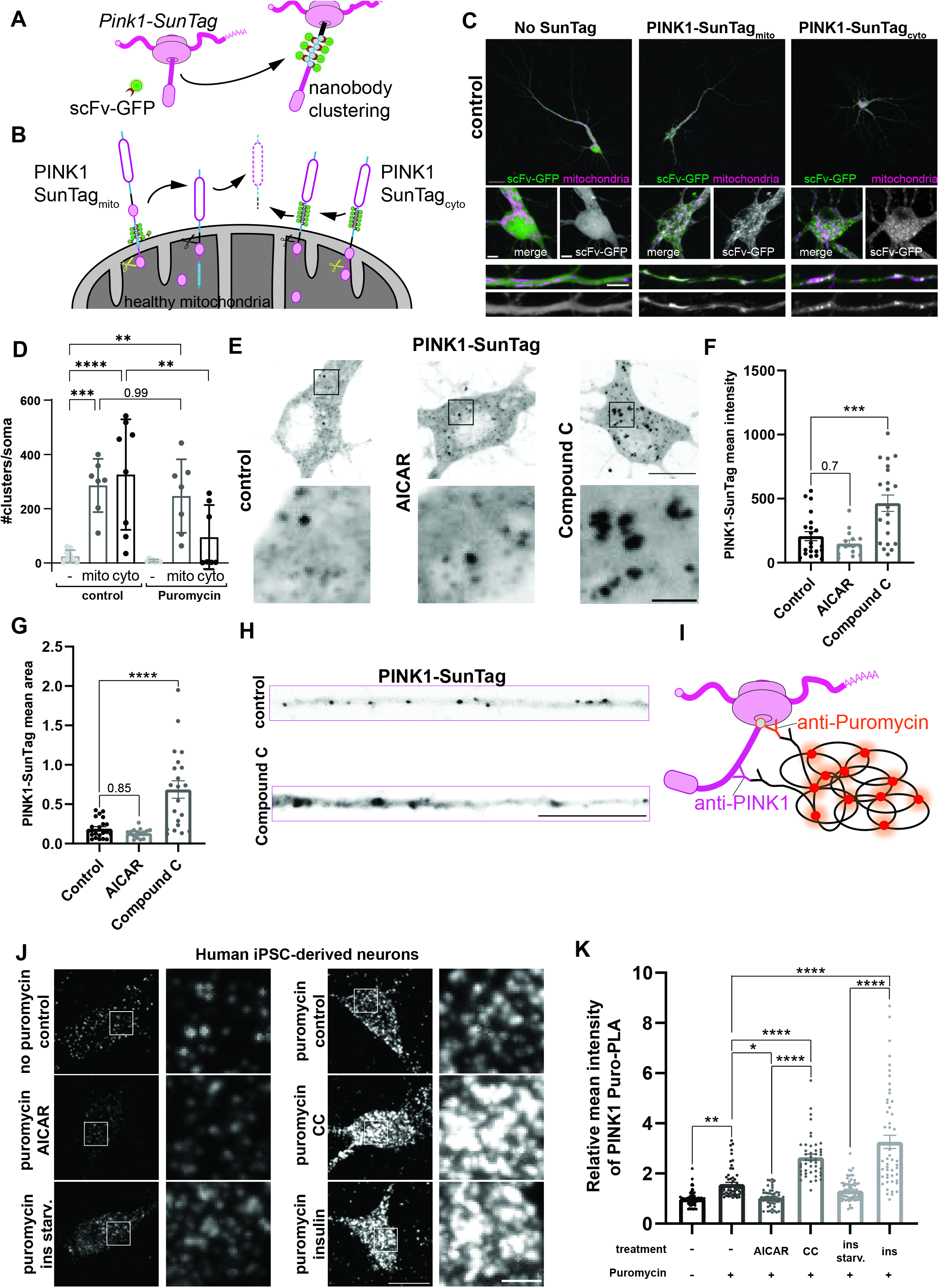
Visualization and regulation of PINK1 translation. **A** Schematic depiction of the SunTag assay. **B** Comparison of the import and processing patterns expected upon insertion of the SunTag repeats within the PINK1 protein for PINK1-SunTagmito and PINK1-SunTagcyto reporters. Dashed lines indicate proteasomal degradation. Scissors represent proteolytic cleavage. **C** Representative images showing the distribution of either the nanobody scFv-GFP alone or cotransfected with either PINK1-SunTagmito or PINK1-SunTagcyto reporters in mouse hippocampal neurons (green). Mitochondria are visualized by coexpression with Cox8-BFP protein (magenta). Inserts showing magnified somas and dendrites are included below. Black-white images correspond to the scFv-GFP signal only. **D** Quantification of the number of clusters per soma following control or puromycin (100 µg/ml, 30 min) treatment as in **C** and **S1B**. One-way ANOVA followed by Tukey’s post hoc test; n = 9-10; p < 0.01 (**), p < 0.001 (***), p < 0.0001 (****). **E** Representative images of cytosolic PINK1 protein clusters visualized by the SunTag system upon AMPK activation or inhibition using AICAR (1 mM, 2 h) or CC (20 µM, 2 h), respectively, in mouse hippocampal neurons. **F** Quantification of the PINK1 protein cluster mean intensity as in **E**. One-way ANOVA followed by Tukey’s post hoc test; n = 15-22; p < 0.001 (***). **G** Quantification of the PINK1 protein cluster mean area as in **E**. One- way ANOVA followed by Tukey’s post hoc test; n = 15-22; p < 0.0001 (****). **H** Representative images of cytosolic PINK1 protein clusters visualized by the SunTag system upon control (untreated) or CC (20 µM, 2 h) treatment in neurites of hippocampal neurons. **I** Schematic illustrating the proximity ligation assay coupled with puromycylation (Puro-PLA) for PINK1. **J** Representative images of PINK1-Puro-PLA signal in human iPSC-derived neurons either untreated (control) or treated with AICAR (1 mM, 2 h), CC (20 µM, 2 h), insulin (500 nM, 1 h) or in insulin-free medium (ins. starv., 2 h), combined with or without Puromycin treatment as indicated (10 µg/ml, 5 min). **K** Quantification of the mean intensity of the PINK1-Puro PLA signal as in **J**. One-way ANOVA followed by Tukey’s post hoc test; n = 11-39; p < 0.05 (*), p < 0.01 (**), p < 0.0001 (****). Data are expressed as mean±SEM. All data points correspond to single cells coming from ≥3 biological replicates (**D,F,G,K**). Scale bars, 25 µm for whole neurons images (**C**) or 5 µm for somas and dendrites (**C**), 10 µm for whole soma and neurites (**E,H,J**), 2 µm for insets (**E,J**).

Treatment with the AMPK inhibitor Compound C (CC) dramatically increased the size and intensity of the SunTag clusters in neuronal cell bodies and neurites (Fig. 1E-H) but not in HeLa cells (Fig. S2A-C), consistent with the neuron-specific mRNA tethering and a prolonged cytosolic dwell-time of the PINK1 precursor (*13*, *14*). Treatment with the AMPK Activator AICAR had little effect (Fig. 1E-H, consistent with already high AMPK activity in the imaging medium lacking insulin (*14*). The effect of CC was not due to altered PINK1 processing (Fig. S2D) or unspecific clustering of the SunTag nanobody (Fig. S2E). The increased signal could be prevented by inhibition of protein translation (Puromycin, Fig. S2F-H). Insulin treatment also induced increased PINK1-SunTag clusters that were reduced by either inhibition of insulin receptor downstream signalling (Wortmannin, Fig. S2I-K) or expression of a phospho-mimetic version of SYNJ2BP (S21E, Fig. S2L-M) that prevents the untethering of the *Pink1* mRNA (*14*). Within these enlarged SunTag clusters we confirmed the simultaneous presence of both *Pink1* mRNA as well as ribosomes, either using MS2/PP7 split-Venus imaging of the same construct (*13*, *14*) (Fig. S3A-B) or a proximity ligation assay (PLA) between an antibody against the small ribosomal protein Rps6 and the Fc portion of the scFv-GFP nanobody used for the SunTag visualization (Fig. S3C-D), respectively. Omission of either the primary anti-Rps6 antibody (Fig. S3E) or the PINK1-SunTag reporter required to cluster the scFv-GFP nanobodies (Fig. S3F) served as controls for the specificity of the PLA signal. PINK1 SunTag clusters almost ubiquitously co-localized with the *Pink1* mRNA and the PLA signal, indicative of active engagement of the mRNA with translating ribosomes.

To assay endogenous PINK1 translation, we used an anti-human PINK1-specific antibody to perform PLA after puromycylation (Puro-PLA) in iPSC-derived human cortical neurons (*18*). After a short pulse of a low Puromycin dose, coincidence-detection of Puromycin with PINK1 by PLA signal (using anti-Puromycin and anti-PINK1 antibodies) allowed the detection of puromycylated PINK1 nascent chains produced within this time interval (Fig. 1I). We confirmed enhanced PINK1 translation upon AMPK inhibition by either CC or insulin treatment (Fig. 1J- K). In contrast, activation of AMPK by either AICAR treatment (1 mM, 2 h) or insulin withdrawal for two hours reduced the observable PLA intensity to background levels (without Puromycin control, Fig. 1J-K). These observations are consistent with the interpretation that the increased size and intensity of the PINK1-SunTag reporter clusters in mouse hippocampal neurons upon AMPK inhibition also represent an increase in overall PINK1 translation and not increased protein stability. Indeed, the degradation kinetics of PINK1 remained unchanged (Fig. S3G-H). Thus, the untethering of the *Pink1* mRNA alters its ribosomal association and increases translation.

## Localization of PINK1 translation at endolysosomes upon AMPK inhibition

As Rab7-positive late endosomes have previously been suggested to form translational hotspots for nuclear-encoded mitochondrial transcripts in neurons (*19*), we tested whether the colocalization of PINK1-SunTag clusters with either mitochondria or late endosomes was affected by AMPK inhibition. Treatment with CC induced a significant shift in the colocalization of the PINK1-SunTag reporter with mitochondria towards Rab7-positive late endosomes, and even stronger towards endolysosomes (TMEM192) or lysosomes (LAMP1) (Fig. 2A-C). This effect was independent of the morphological changes in the PINK1 clusters, as there was no significant change in the colocalization coefficients with any of the organellar markers upon rotation of one of the channels prior to analysis (Fig. S4A-D). The shift in localization of the PINK1 preprotein was mirrored by the distribution of its mRNA, which was also increasingly localized to endolysosomes following CC treatment (Fig. S4E-F) (*14*), and this could be prevented by forced mRNA tethering to mitochondria via expression of phospho-mimetic SYNJ2BP-S21E (Fig. S4G-I), analogous to the reduction in PINK1-SunTag intensity (compare to Fig. S2L-M analyzing the same set of neurons). However, some mitochondrial localization remained, placing the PINK1 sites of translation at organellar contact sites (Fig. 2D-E). This formation of organellar contact sites may be necessary to ensure later targeting of the PINK1 preprotein to mitochondria.

**Fig. 2.**
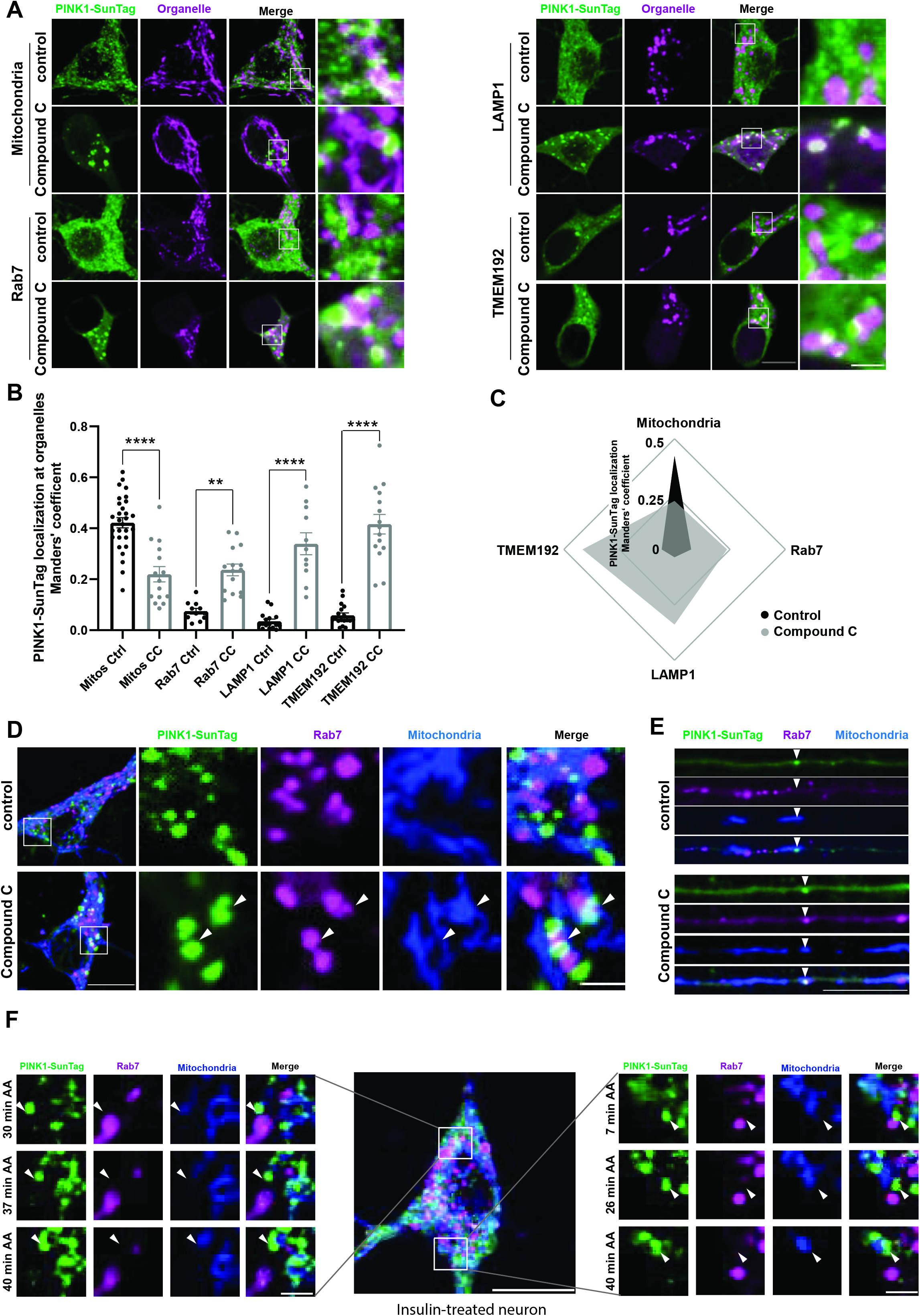
Localization of PINK1 translation at endolysosomes upon AMPK inhibition. **A** Representative images of cytosolic PINK1 protein clusters visualized by the SunTag system and fluorescently-labeled organelles (Cox8-mRaspberry (mitochondria), Rab7-mCherry, LAMP1-mCherry or TMEM192-RFP) upon control or CC (20 µM, 2 h) treatment in hippocampal neurons. **B** Quantification of the Manders’ coefficient of the colocalization between the PINK1-SunTag signal and the respective organellar signal as in **A**. One-way ANOVA followed by Tukey’s post hoc test; n = 11-29; p < 0.01 (**), p < 0.0001 (****). **C** Radar chart displaying the colocalization of PINK1-SunTag clusters with different organelles (mitochondria, Rab7-, LAMP1-, and TMEM192-positive endolysosomes) upon control or CC (20 µM, 2 h) treatment as in **B**. **D** Representative images of cytosolic PINK1 protein clusters visualized by the SunTag system and fluorescently-labeled organelles (mito-BFP and Rab7-mCherry) upon control or CC (20 µM, 2 h) treatment in the soma of hippocampal neurons. Arrowheads point to regions where PINK1 SunTag clusters, late endosomes and mitochondria are in close proximity. **E** Representative images of cytosolic PINK1 protein clusters visualized by the SunTag system and fluorescently-labeled organelles (mito-BFP and Rab7-mCherry) upon control (untreated) or CC (20 µM, 2 h) treatment in a neurite of hippocampal neurons. Arrowheads point to PINK1 SunTag clusters at mitochondria without (control) or with (CC) a late endosome in proximity. **F** Representative images of cytosolic PINK1 protein clusters visualized by the SunTag system and fluorescently-labeled organelles (mito-BFP and Rab7-mCherry) in an insulin (500 nM, 1 h)- treated hippocampal neuron following 20 µM AA treatment. Arrowheads point to PINK1 SunTag clusters at mitochondria-late endolysosome contact sites moving away from the endolysosome and towards mitochondria. Data are expressed as mean±SEM. All data points correspond to single cells coming from ≥3 biological replicates (**B**). Scale bars, 10 µm for whole soma, 2 µm for insets.

To visualize the journey of the PINK1 protein from its site of synthesis towards mitochondria, we performed live cell imaging of neurons after induction of PINK1 synthesis with insulin for 1 h, and induced mitochondrial damage with Antimycin A (AA, 20 µM). We could observe that the PINK1-SunTag signal located at mitochondria-endolysosomal contact sites moved away from the endolysosome and towards mitochondria (Fig. 2F left and Supplemental movie 1), sometimes in concert with mitochondrial shrinkage typical of mitochondrial damage (Fig. 2F right and Supplemental movie 2). Finally, damaged mitochondria appear surrounded by a halo of stabilized PINK1 (Fig. S5A-C).

## ER tubules contribute to PINK1 biogenesis

To observe the mitochondria-endolysosomal contact sites in more detail, we performed correlative light and electron microscopy (CLEM) in transfected neurons upon CC treatment (Fig. 3A-B). CLEM images confirmed that the large PINK1-SunTag clusters were indeed in close apposition to endolysosomes and mitochondria (Fig. 3A). Surprisingly, we also identified ER tubules in-between mitochondria and endolysosomes as well as directly overlapping the PINK1-SunTag signal in such clusters (Fig. 3B inset, green arrowheads). This suggested that instead of a direct mitochondria-endolysosomal contact site, the ER may bridge the two organelles, as well as be engaged in PINK1 translation.

**Fig. 3.**
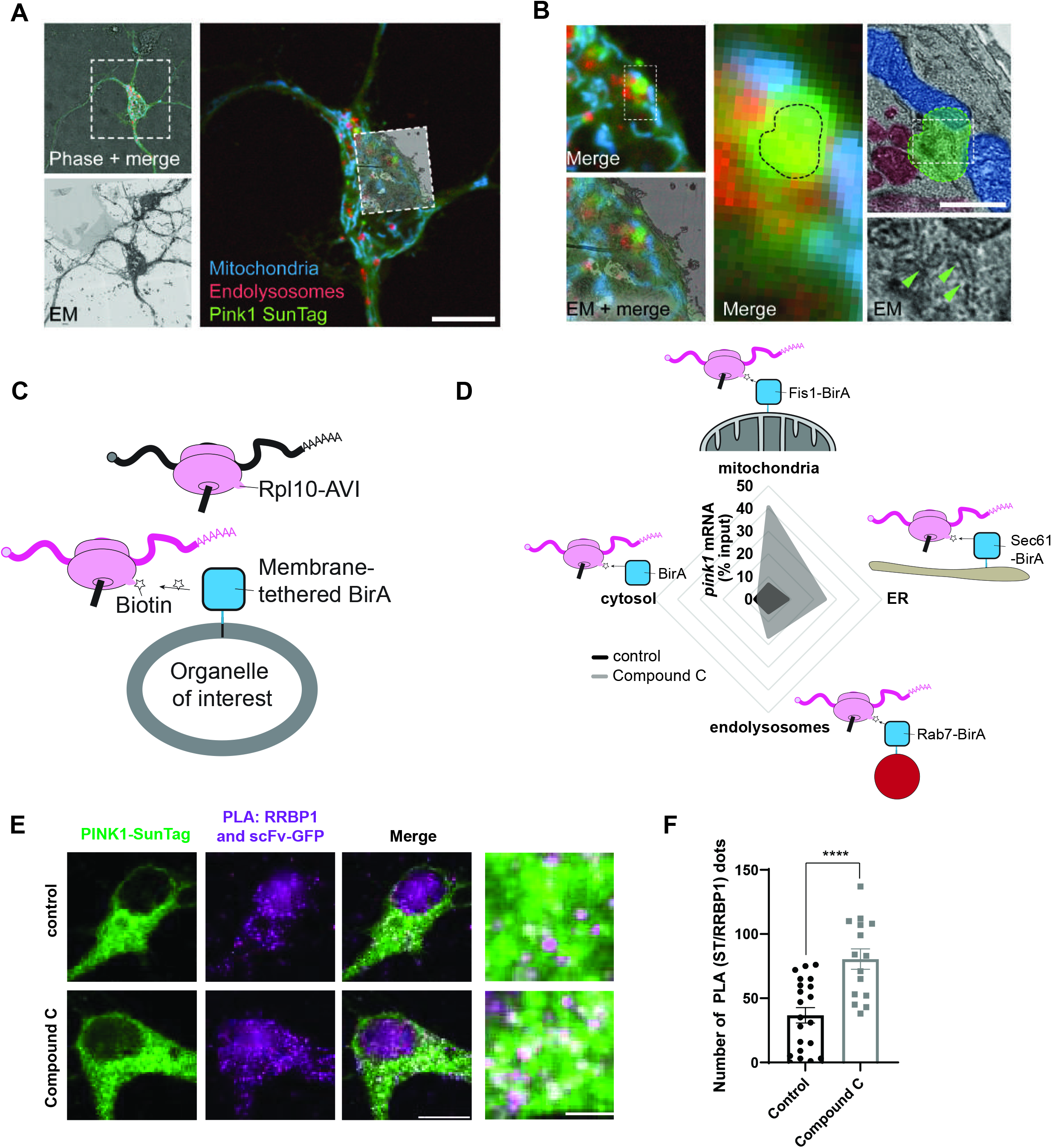
ER tubules contribute to PINK1 biogenesis. **A** Correlated light and electron microscopy (CLEM) of CC treated mouse hippocampal neurons, showing the correlation between phase, fluorescence and electron microscopy images at the cellular level. **B** Correlated light and electron microscopy (CLEM) as in **A** at the subcellular level. A PINK1-SunTag cluster (green) is located where two mitochondria (cyan) are in close proximity to a cluster of endolysosomes (red), and the electron micrograph (lower right inset) reveals an ER tubule within the area covered by the PINK1-SunTag signal (green arrowheads). Boxes indicate the location of insets. **C** Schematic representation of the BirA assay. BirA is targeted to different organelles. BirA selectively biotinylates its substrate AVI-Tag, which is fused to the ribosomal protein Rpl10 (Rpl10-AVI). **D** Radar chart displaying the association of endogenous *Pink1* mRNA (relative to the input) with BirA-biotinylated ribosomes in the cytosol or in proximity to different organelles (mitochondria, ER, endolysosomes) in mouse cortical neurons, upon control (untreated) or CC (20 µM, 2 h) treatment. Values represent the average of 5 independent experiments. **E** Representative images of cytosolic PINK1 clusters visualized by the SunTag system as well as the PLA signal for scFv-GFP and RRBP1 upon control and CC (20 µM, 2 h) treatment in hippocampal neurons. **F** Quantification of the number of PLA clusters per soma as in **E**. Student’s t-test; n = 15-21; p < 0.0001 (****). Data are expressed as mean±SEM (**F**) or mean (**D**). Data points correspond to single cells coming from ≥3 biological replicates (**F**). Scale bars, 10 µm for CLEM (**A**) and whole soma (**E**), 1 µm for inset in **B**, 2 µm for inset in **E**.

The ER is well known for its role as a major hub for protein translation. To evaluate the subcellular location of the ribosomes translating *Pink1* mRNA we performed proximity biotinylation of ribosomes at different organellar locations. Here, the specific biotinylation of an AVI-tag fused to the ribosomal subunit Rpl10a (Rpl10-AVI) by the AVI-specific BirA Ligase targeted to an organellar membrane of interest (*20*) was used to label the specific population of ribosomes associated with this subcellular location (Fig. 3C). As expected, we observed an increased association of the *Pink1* transcript with membrane-associated ribosomes upon inhibition of AMPK, while the fraction of *Pink1* in cytoplasmic ribosomes slightly reduced (Fig. 3D). This is in contrast to a control transcript (*actin*), whose distribution among membrane-associated ribosomes was unresponsive to AMPK inhibition (Fig. S6A-B). This is consistent with the idea that membrane-associated ribosomes at ER-endolysosome-mitochondria contact sites are the main source of PINK1 preprotein upon *Pink1* mRNA untethering.

As SYNJ2BP in other cell types interacts with the ribosome binding protein 1 (RRBP1, also known as p180), an ER-localized protein that forms a contact site rich in ribosomes (*21–23*), we tested whether RRBP1 would also play a role in the biogenesis of PINK1. Consistent with the increased ER-localized translation of PINK1, PLA experiments revealed an increase in the proximity between RRBP1 and the PINK1-SunTag reporter upon CC treatment (Fig. 3E-F). This was specific to RRBP1, as no difference was observed with Sec61, another ribosomal receptor on the ER (Fig. S6C-D). However, silencing of RRBP1 by shRNA (Fig. S7A-B) neither altered the localization of the large CC-induced PINK1-SunTag clusters (Fig. S7C-E), nor the ability of PINK1 to induce mitophagy by phosphorylation of ubiquitin in response to mitochondrial damage (Fig. S7F-G). These data suggest that while RRBP1 may be involved in the ER-associated translation of PINK1, removing it by knock down was not sufficient to disrupt the mitochondria-ER tether necessary for targeting the PINK1 precursor to mitochondria.

## PINK1 import follows the ER-SURF pathway

In yeast, the ER has recently been shown to facilitate the targeting of mitochondrial precursor proteins. This occurs through ER membrane-associated chaperones, including Djp1, which guide unfolded preproteins to the mitochondrial import pores in a pathway termed ER-SURF (*24*, *25*). We therefore tested, whether DNAJB6, the mammalian homolog of Djp1, colocalized with the PINK1-SunTag clusters induced by CC treatment. Indeed, we observed a striking colocalization in clusters between this chaperone and the PINK1-SunTag reporter upon CC treatment (Fig. 4A). Importantly, this was not driven by overexpression of PINK1 or the SunTag reporter system, as clustering of endogenous DNAJB6 could be observed upon CC treatment in untransfected neurons (Fig. 4B-C) but not in HeLa cells (Fig. S8A). Knock-down of DNAJB6 by shRNA (Fig. S8B-C and (*26*)) did not prevent the formation or the localization of the large PINK1-SunTag clusters in response to CC (Fig. S8D-F), consistent with the role of the chaperone acting downstream of increased PINK1 translation.

**Fig. 4.**
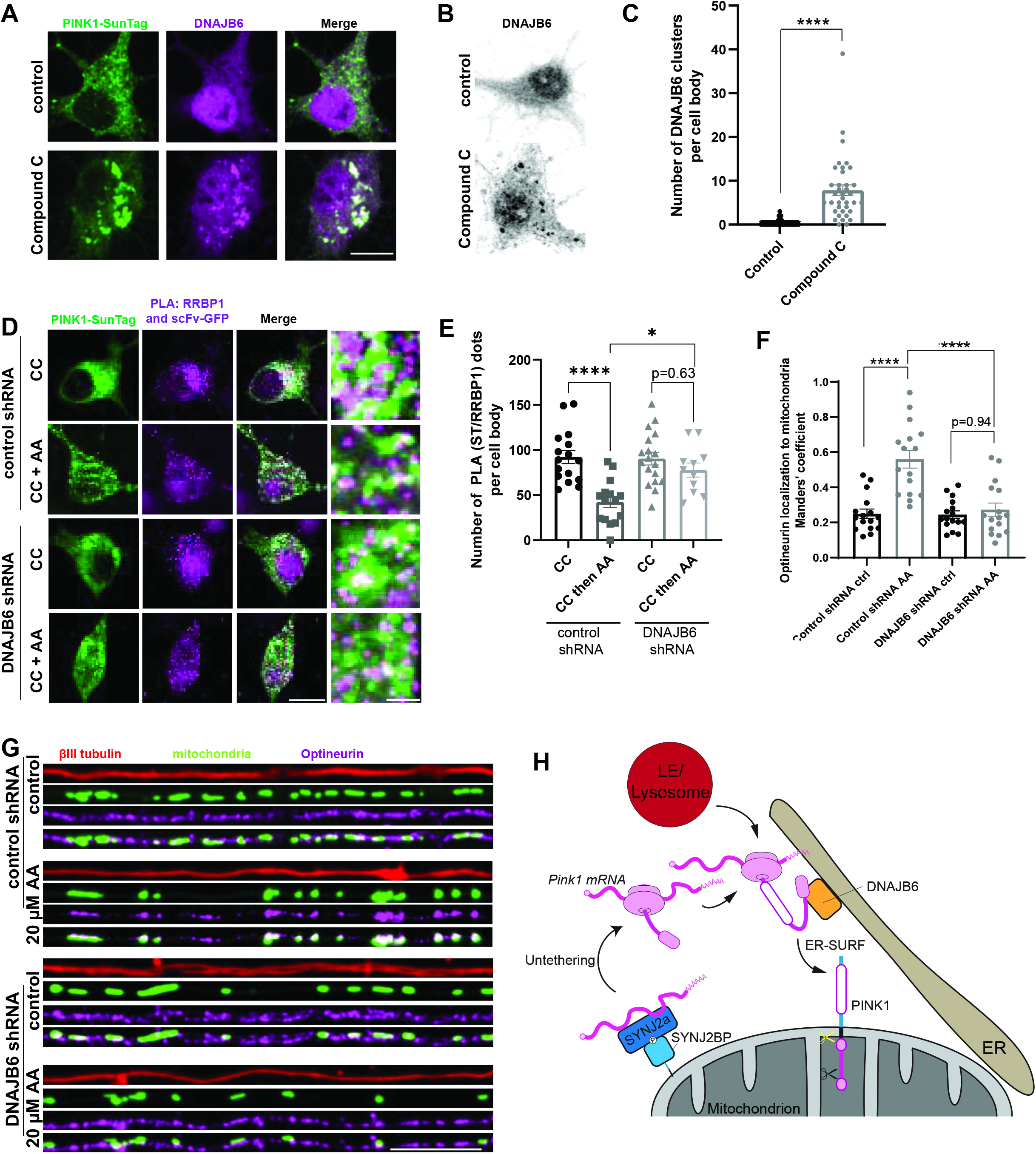
PINK1 import follows the ER-SURF pathway. **A** Representative images of cytosolic PINK1 protein clusters visualized by the SunTag system and immunostaining against endogenous DNAJB6 upon control (untreated) or CC (20 µM, 2 h) treatment in hippocampal neurons. **B** Representative images of DNAJB6 immunostaining upon control (untreated) and CC (20 µM, 2 h) treatment in hippocampal neurons. **C** Quantification of the number of DNAJB6 clusters per cell body as in **B**. Student’s t-test; n = 37-44; p < 0.0001 (****). **D** Representative images of cytosolic PINK1 clusters visualized by the SunTag system as well as the PLA signal for scFv-GFP and RRBP1 upon CC (20 µM, 2 h) and AA (20 µM, 45 min) treatment in hippocampal neurons that express either control or DNAJB6 shRNA. **E** Quantification of the number of PLA clusters per soma as in **D**. One-way ANOVA followed by Tukey’s post hoc test; n = 15-18; p < 0.05 (*), p < 0.0001 (****). **F** Quantification of the Manders’ coefficient for colocalization of the mitochondrial and the Optineurin signal in neurites expressing control shRNA or DNAJB6 shRNA treated with vehicle (control) or AA (20 µM, 45 min). One- way ANOVA followed by Tukey’s post hoc test; n = 16; p < 0.0001 (****). **G** Representative images of neurites of hippocampal neurons overexpressing mito-meGFP as well as either control shRNA or DNAJB6 shRNA. The neurons were treated with vehicle (control) or AA (20 µM, 45 min) and immunostained against optineurin and βIII tubulin. **H** Schematic illustrating PINK1 biogenesis in neurons involving *Pink1* mRNA untethering from mitochondria, translation in proximity to functional endolysosomes and ER, and DNAJB6-mediated ER-SURF guarding the PINK1 protein during its journey towards its mitochondrial destination. Data are expressed as mean±SEM. All data points correspond to single cells coming from ≥3 biological replicates (**C,E,F**). Scale bars, 10 µm.

In contrast, silencing of DNAJB6 prevented the association and stabilization of PINK1 in response to mitochondrial damage (Fig. S9A-F). Instead of translocating to mitochondria, the PINK1 preprotein remained associated with endolysosomes upon CC treatment in the absence of DNAJB6 (Fig. S9A-C). To observe the dynamics of the PINK1 precursor, we imaged the same cell before and after addition of Antimycin A in the presence of insulin as a more physiological AMPK inhibitor. Again, the PINK1 reporter failed to move to mitochondria (Fig. S9D-E).

Instead, distinct PINK1 clusters remained associated with Rab7-positive endolysosomes (Fig. S9F), arguing that the lack of DNAJB6 prevented their transfer to mitochondria Moreover, upon DNAJB6 silencing, the association PINK1-SunTag clusters with RRBP1 was not responsive to mitochondrial depolarization (Fig. 4D-E), suggesting that translated PINK1 was not efficiently transferred to mitochondria and remained close to the ER. Knock down of DNAJB6 also prevented effective mitophagy in neurites, as the recruitment of the autophagy adaptor Optineurin (Fig. 4F-G) and the colocalization of mitochondrial material with lysosomes after AA treatment were significantly reduced (Fig. S10A-B). This block in PINK1 activity despite ongoing protein translation was consistent with the ER-SURF model for PINK1 biogenesis, in which the PINK1 protein is translated on ribosomes located in proximity to the ER and endolysosomes, before being transferred to the mitochondrial import pore via association with DNAJB6 at the ER surface (Fig. 4H). Together, AMPK inhibition induces the relocation of PINK1 synthesis away from mitochondria. For proper PINK1 synthesis and activity at mitochondria the PINK1 precursor is then shuttled along the ER from its site of synthesis to the site of protein import at mitochondria with the help of the ER-associated chaperone DNAJB6.

## Discussion

Translation of proteins in highly polarized cells such as neurons requires a concerted action of several organelles. We and others have shown that hitch-hiking of mRNAs occurs on several organelles, including mitochondria, early endosomes, late endosomes, and lysosomes (*13*, *19*, *27–29*). Likewise, ribosomes are transported into axons in association with endosomes and the ER (*23*, *30*). However, the functional connection of these organelles to the process of local translation remains unclear. Using a live-cell imaging tool for the mitochondrial quality control protein PINK1, we now uncover that the ER bridges the respective organelles and allows the PINK1 preprotein to follow the ER-SURF pathway for mitochondrial protein import.

Unlike the ER, the mitochondrial import pores can import proteins in a post-translational manner, but no mechanisms are known that would prevent the premature association of the nascent chain with the receptors of the TOM complex. Instead, mechanisms anchoring the ribosome and/or mRNAs to the mitochondrion have been described, which rather support co-translational import (*31–33*). However, constant mitochondrial association of *Pink1* mRNA prevents PINK1 function in neurons (*14*), arguing that the mRNA needs to be untethered to produce a fully functional protein. Untethering of *Pink1* mRNA might be a prerequisite for mitophagy as it increases the cytosolic dwell time and potentially inhibits co-translational import of the PINK1 precursor. This may facilitate the interaction between N- and C-terminal parts of PINK1 post-translationally, which is necessary for the arrest of PINK1 at the TOM complex upon mitochondrial damage (*34*). The close apposition of endolysosomes may hereby ensure timely degradation of the mitophagosome.

Intriguingly, PINK1 may also be produced at endomembrane-localized ribosomes in other cell types, given that small amounts of *Pink1* mRNA ribosomal footprints were recovered with ER-associated ribosomes in HEK293 cells unlike most transcripts encoding mitochondrially targeted proteins (*20*). PINK1 maturation has already been suggested to involve mitochondria-ER contact sites and PINK1 mitochondrial levels are regulated by the ER-associated degradation (ERAD) machinery and especially the ATPase Valosin-containing protein (VCP) (*35*, *36*). In that context, the interaction with the ER ensures the timely degradation of the PINK1 protein after its processing in mitochondria. Our data now suggest that the ER may serve a dual role by producing the preprotein on membrane-associated ribosomes and by providing part of the degradative machinery to regulate timely destruction of the PINK1 protein to suppress mitophagy in the absence of mitochondrial damage. In other cell types, PINK1 may not exhibit a comparable increase in ER-associated translation upon AMPK inhibition as in neurons, probably due to the differential expression of the *Pink1* mRNA RBP SYNJ2a (*13*). Nonetheless, this does not rule out the possibility that, in non-neuronal cells, the ER could be the default site for PINK1 synthesis, due to the lack of *Pink1* mRNA tethering complex and hence an absence of regulation by AMPK and insulin signalling.

The ER also supplies chaperones to safeguard the PINK1 preproteins from aggregation on its route to mitochondria, analogous to the ER-SURF pathway observed in yeast (*24*). We have confirmed an essential role for DNAJB6, the mammalian homolog of the ER-SURF chaperone Djp1, for PINK1 function in neurons. Transfer of the ER-localized PINK1 precursor presumably occurs at a mitochondria-ER contact site. We observe the PINK1 precursor in close proximity with RRBP1 upon untethering of the SYNJ2a-SYNJ2BP interaction, yet knock down of RRBP1 did not prevent PINK1 action at mitochondria, suggesting that other mitochondria-ER contact sites can substitute for the loss of RRBP1-SYNJ2BP interaction. This is in line with the current model of the ER-SURF pathway in yeast, in which loss of multiple mitochondria-ER contact sites is necessary to prevent mitochondrial import via the ER-SURF pathway (*25*).

DNAJB6 has been previously implicated in PINK1/Parkin-mediated mitophagy, demonstrated by its ability to overcome the failure of a Parkin RING domain mutant (C289G). This rescue is likely attributed to its function as a disaggregase for this aggregation-prone Parkin mutant (*37*). Similarly, its overexpression can reduce the toxicity of other aggregation-prone proteins, including alpha-synuclein or Huntingtin (*38*, *39*) and DNAJB6 is even found at the core of Lewy bodies, the pathological hallmark of PD (*40*). Our identification of DNAJB6 as a crucial factor in PINK1 activation provides a connection between these aggregates and mitochondrial quality control. It is conceivable that sequestration of DNAJB6 into Lewy bodies might diminish the available pool for ER-SURF of the PINK1 protein precursor, consequently affecting mitochondrial quality control. As our results are mainly derived from *in vitro* cultured mouse neurons that do not develop Lewy body pathologies, future studies will be needed to confirm this hypothesis. By elucidating the locally active mechanisms governing mitochondrial protein translation and targeting, novel approaches to improve mitochondrial biogenesis and quality control could emerge.

## Materials and methods Cell culture preparation

### Mouse primary hippocampal and cortical neurons

Mouse primary hippocampal and cortical neurons were obtained from embryonic day 16.5 (E16.5) embryos of timed-pregnant mice as previously described (13,14). The mouse procedures in this study received approval from the Government of Upper Bavaria and were conducted in accordance with relevant guidelines and regulations. C57BL/6 WT mice were housed in the animal facility of the Max Planck Institute for Biological Intelligence, Martinsried, Germany. The animals were provided with a controlled environment including unrestricted access to food and water. Experiments involved animals of both sexes.

The timed-pregnant mice were euthanized using CO2 followed by the rapid extraction of the E16.5 embryos from the uterus. The embryos were placed into ice-cold dissociation medium (Ca^2+^-free Hank’s Balanced Salt Solution with 100 mM MgCl2, 10 mM kynurenic acid, and 100 mM HEPES). After removal from the amniotic sac, the embryos were immediately decapitated using forceps. Under a dissecting microscope, the brains were extracted from the skull followed by removal of the cerebellum, the midbrain and the meninges resulting in the isolation of the two cortices. From each hemisphere, the hippocampus was dissected, and the collected hippocampi and cortices were placed in distinct tubes containing ice-cold dissociation medium. To enzymatically dissociate the tissue, papain/L-cysteine (Sigma-Aldrich) was applied and after a 5- min incubation at 37 °C stopped by addition of trypsin inhibitor (Abnova). Subsequently, the tissue underwent three washes with NB+B27+PSG medium [Neurobasal medium containing B27, 500 µg/ml L-glutamine, and 100 U/ml penicillin/100 µg/ml streptomycin (1 % P/S) (all from Thermo Fisher Scientific)]. Finally, the hippocampal and cortical tissues were separately dissociated into single cells through trituration (10-15 times) using a p1000 pipette-tip. Hippocampal and cortical neurons were seeded on 24-well glass bottom plates (CellVis), on acid-washed glass coverslips (1.5 mm, Marienfeld) or on 6-well plates (Greiner), which were previously coated with 3.5 µg/ml laminin (Thermo Fisher Scientific) and 20 µg/ml poly-L-lysine (PLL; Sigma-Aldrich), and cultivated in NB+B27+PSG. The primary neurons were maintained in a humidified incubator with 5 % CO2 at 37 °C. Half of the medium was replaced with fresh NB+B27+PSG every three to five days.

## Human induced pluripotent stem cell (iPSC)-derived cortical neurons

Human induced pluripotent stem cells (iPSCs; cell line HPSI0314-hoik_1) were acquired from the Wellcome Trust Sanger Institute HipSci Repository. They were cultured in StemFlex medium (Thermo Fisher Scientific) in 10 cm dishes (Falcon) coated with Matrigel (Corning). The StemFlex medium was daily renewed, and iPSCs were passaged at 80 % confluency using the enzyme-free passaging reagent ReLeSR (Stem Cell Technologies).

To induce the differentiation of iPSCs into cortical neurons, the previously described protocol involving the overexpression of the transcription factor neurogenin-2 (NGN-2) was used (14, 43). Briefly, on day -2, iPSCs were dissociated into single cells using Accutase (Thermo Fisher Scientific) and seeded in 10 cm dishes coated with Matrigel at a density of 2.5x10^6^ cells per dish in StemFlex medium containing 10 µM Y-27632 (Tocris). 24 h after seeding, on day -1, the cells underwent lentiviral transduction with FudeltaGW-rtTA and pLV-TetO-hNGN2-eGFP-puro in StemFlex medium, which are essential for subsequent differentiation into cortical neurons. Following a 5-h incubation, the medium was exchanged with fresh StemFlex supplemented with 2 µg/ml doxycycline (Takara) to initiate TetO gene expression. On day 0, the cells were exposed to N2/DMEM/F12/NEAA (N2 medium; Thermo Fisher Scientific) containing 2 µg/ml doxycycline, 10 ng/ml NT-3 (PeproTech), 0.2 µg/ml laminin (Thermo Fisher Scientific) and 10 ng/ml BDNF (PeproTech). On day 1, a 24-h Puromycin selection phase was initiated by introducing fresh N2 medium containing 1 µg/ml Puromycin (Enzo Life Sciences), 2 µg/ml doxycycline, 10 ng/ml NT-3, 0.2 µg/ml laminin, and 10 ng/ml BDNF to the cells. On day 2, the transition from N2 medium to B27/Neurobasal-A/Glutamax (B27 medium; Thermo Fisher Scientific) took place, supplemented with 2 µg/ml doxycycline, 10 ng/ml NT-3, 0.2 µg/ml laminin, 10 ng/ml BDNF, and 2 µM Ara-C (Sigma-Aldrich). From day 3 to 6, medium replacement occurred every other day (day 3 and 5) using fresh B27 medium supplemented with 2 µg/ml doxycycline, 10 ng/ml NT-3, 0.2 µg/ml laminin, 10 ng/ml BDNF, and 2 µM Ara-C. On day 7, the B27 medium was replaced by conditioned Südhof neuronal growth medium (NGN2 glial conditioned medium/B27/Neurobasal-A/NaHCO3/Glucose/Transferrin/L-Glutamine) containing 10 ng/ml NT-3, 0.2 µg/ml laminin, and 10 ng/ml BDNF. Additionally, the medium was supplemented with 10 µM Y-27632 to prime the cells for re-plating. The re-plating of iPSC-derived cortical neurons took place on day 8, involving cell dissociation using TrypLE Express (Thermo Fisher Scientific) for 5 min at 37 °C. Following enzyme inactivation using fetal bovine serum (FBS; Thermo Fisher Scientific), a 5-min centrifugation at 200 x g, and resuspension in fresh conditioned Südhof neuronal growth medium containing 10 ng/ml NT-3, 0.2 µg/ml laminin and 10 ng/ml BDNF, the cells were filtered through a 70 µm cell strainer. Subsequently, the cells were seeded on 6-well plates (Greiner) or on acid-washed glass coverslips (1.5 mm, Marienfeld) at a density of 2x10^6^ or 10^5^ cells per well, respectively. Human iPSC-derived neurons were maintained in a humidified incubator with 5 % CO2 at 37 °C. Half of the medium was replaced with fresh conditioned Südhof neuronal growth medium supplemented with 10 ng/ml NT-3, 0.2 µg/ml laminin and 10 ng/ml BDNF every other day.

## HEK293T and HeLa cells

Human embryonic kidney 293-T (HEK293T) and Henrietta Lacks cervical carcinoma (HeLa) cells were obtained from ATCC®. They were cultured in T75 flasks (Falcon) using Dulbecco’s modified Eagle medium (DMEM) containing Glutamax (Thermo Fisher Scientific), 10 % FBS and 1 % P/S. HEK293T and HeLa cells were maintained in a humidified incubator at 37 °C with 5 % CO2. Passaging of cells was performed twice a week at approximately 80 % confluency utilizing Trypsin-EDTA (Thermo Fisher Scientific). Experiments were performed until cells reached passage 15-20.

## DNA constructs

Mito-meGFP, mito-mRaspberry-7, mito-BFP, mCherry-Rab7A, pLAMP1-mCherry, TMEM192- mRFP-3xHA, nls-ha-MCP-VenusN-nls-ha-PCP-VenusC, scFv-GCN4-sfGFP (scFv-sfGFP), and AviTag-RPL10a were purchased from Addgene (#172481, #55931, #49151, #61804, #45147, #134631, #52985, #60907 and #62365 respectively). The plasmids required for neuronal differentiation of iPSCs FudeltaGW-rtTA and pLV-TetO-hNGN2-eGFP-puro as well as the plasmids for virus production pMDLg/pRRE, pMD2.G and pRSV-Rev were acquired from Addgene as well (#19780, #79823, #12251, #12259 and #12253, respectively). A non-targeting control shRNA plasmid (TR30021) as well as the plasmids coding for an shRNA against DNAJB6 (TRCN0000139049) or RRBP1 (TRCN0000194406) in a pLKO vector backbone were purchased from Sigma-Aldrich.

Pink1-kinase dead-MS2-PP7 and myc-tagged SYNJ2BP-WT (*13*) as well as myc-tagged SYNJ2BP-S21E (*14*) plasmids have been previously described. The Pink1-kinase dead- 10xGCN4mito-12xMS2-PP7 (Pink1-SunTagmito) and Pink1-kinase dead-10xGCN4cyto-12xMS2- PP7 (Pink1-SunTagcyto) were generated by inserting 10x GCN4 consecutive repeats (obtained by enzymatic restriction) in frame between the amino acid positions 50-56 or 141-145 of rat PINK1, respectively. Compatible target positions were generated at the indicated positions by site directed mutagenesis by PCR using Q5 High-Fidelity DNA Polymerase (NEB). Constructs were grown on Stbl3 chemically competent *E. coli* bacteria (Thermo Fisher Scientific). The scFv-GCN4- mRaspberry (scFv-mRaspberry) was generated by substituting the sfGFP coding sequence of the scFv-GCN4-sfGFP plasmid by the mRaspberry coding sequence (obtained from mito-mRaspberry-7 plasmid). Both plasmids were enzymatically digested and corresponding fragments were ligated by T4 ligase activity (NEB). The BirA-mCherry-Sec61β construct was generated by PCR amplification of the BirA-mCherry-Sec61β including the Tet-O inducible promoter from the Addgene plasmid #62366 followed by its insertion into the lentiviral vector backbone pMK1047 of the Addgene construct #62365 using the restriction enzymes NheI and EcoRI. The BirA-mCherry-Fis1 construct was generated by first replacing Sec61β coding sequence in the Addgene construct #62366 by the C-terminal part of Fis1 using restriction-free cloning (*41*). Afterwards, BirA-mCherry-Fis1 including the Tet-O inducible promoter was amplified and subsequently inserted into the vector backbone pMK1047 of the Addgene plasmid #62365 as before. The BirA-mCherry-Rab7A construct was generated by amplifying BirA-mCherry including the Tet-O inducible promoter from the Addgene plasmid #62366 and Rab7A from the Addgene plasmid #61804. In a three-way ligation, both inserts were integrated into the vector backbone pMK1047 of the Addgene plasmid #62365 using the restriction enzymes NheI, SbfI, and EcoRI. Cytosolic BirA-mCherry construct was generated by amplifying BirA-mCherry including the Tet-O inducible promoter from the Addgene plasmid #62366, including a stop codon by site directed mutagenesis by PCR. The PCR product was then inserted into the vector backbone pMK1047 of the Addgene plasmid #62365 using the restriction enzymes NheI and EcoRI. Constructs were grown on DH5α chemically competent *E. coli* bacteria (strain Invitrogen, prepared in-house).

## Lentivirus production

Lentiviral particles were generated using HEK293T cells as previously described (*14*). The cells were seeded onto collagen type-I (Sigma-Aldrich)-coated 10 cm dishes (Falcon) at a density of 6x10^6^ cells per dish and cultured in DMEM containing 10 % FBS and 1 % P/S. After 24 h, each 10 cm dish of HEK293T cells underwent transfection with 5 µg of the packaging plasmid mix (pMDLg/pRRE, pRSV-Rev, pMD2.G; ratio 4:1:1) and 5 µg of the relevant transfer plasmid (pLV-TetO-hNGN2-eGFP-puro, FudeltaGW-rtTA, AviTag RPL10a, BirA-mCherry-Fis1, BirA-mCherry-Sec61β, BirA-mCherry-Rab7A, cytosolic BirA-mCherry) using TransIT-Lenti reagent (Mirus Bio). After 48 h, the cell culture medium containing lentiviral particles was harvested, clarified by centrifugation at 300 x g for 10 min at room temperature, and then mixed with lentivirus precipitation solution (Alstem) at a 4:1 ratio. The mixture was incubated at 4 °C overnight before undergoing centrifugation at 1,500 x g for 30 min at 4 °C to precipitate lentiviral particles. After discarding the supernatant, the pellet containing lentiviral particles was reconstituted in 1 ml of ice-cold PBS per 10 cm dish, aliquoted and stored at -80 °C.

## Immunoblotting

For the cycloheximide chase assay, human iPSC-derived neurons were seeded in 6-well plates (Greiner) coated with PLL and laminin at a density of 2x10^6^ cells per well and cultured in conditioned Südhof neuronal growth medium as described above. On day in vitro 14 (DIV14), differentiated neurons were pre-incubated in medium with or without insulin for 2 h followed by 70 µM cycloheximide (Enzo Life Sciences) treatment for 30 min, 60 min or 120 min. Subsequently, the neurons were lysed using a buffer containing 25 mM Tris/HCl pH 7.4, 1 % NP- 40, 150 mM NaCl, supplemented with 1x protease inhibitor cocktail (Roche) and 200 µM PMSF. After centrifugation of the lysates at 9,300 x g for 1 min at 4 °C, the pellet was discarded while the supernatant was mixed with Laemmli sample buffer and boiled at 95 °C for 5 min. Using standard protocols, the samples were electrophoretically separated by SDS-PAGE (12 % polyacrylamide gel) and immunoblotted on Nitrocellulose membranes (Amersham/Cytiva). Membranes were blocked with 1X Blocker FL fluorescent Blocking Buffer (Thermo Fisher Scientific). Antibodies employed for immunoblotting included anti-PINK1 rabbit antibody (1:500; Novus Biologicals, #BC100-494) and anti-βIII tubulin mouse (2G10) antibody (1:2000; Invitrogen, #MA1-118).

For the validation of RRBP1 and DNAJB6 shRNAs, HEK293T cells were seeded in 6-well plates (Greiner) at a density of 0.5x10^6^ cells per well and cultured in DMEM containing 10 % FBS and 1 % P/S. One day after seeding, transfection was performed using calcium phosphate for 5 h introducing control shRNA, shRNA targeting RRBP1 or DNAJB6 into the cells. Three days after transfection, HEK293T cells were lysed using a buffer containing 25 mM Tris/HCl pH 7.4, 1 % NP-40, 150 mM NaCl, supplemented with 1x protease inhibitor cocktail (Roche) and 200 µM PMSF. After centrifugation of the lysates at 9,300 x g for 1 min at 4 °C, the pellet was discarded while the supernatant was mixed with Laemmli sample buffer and boiled at 95 °C for 5 min. Using standard protocols, the samples were electrophoretically separated by SDS-PAGE (12 % polyacrylamide gel) and immunoblotting. Antibodies employed for immunoblotting included anti- RRBP1 rabbit antibody (1:500; Abcam; #ab95983), anti-DNAJB6 rabbit antibody (1:500; Proteintech; #11707-1-AP), and anti-β-actin mouse (AC-74) antibody (1:500; Sigma-Aldrich, #A5316).

For the analysis of the cellular localization and processing of Pink1-SunTagmito and Pink1- SunTagcyto, HEK293T cells were transfected with the indicated plasmids using standard calcium phosphate protocols. After 24-36 h of transfection, cells were incubated with either control medium or medium containing either 100 µM Carbonyl cyanide m-chlorophenyl hydrazone (CCCP; Sigma) or 10 µM MG-132 (Sigma) for 2 h. Then, cells were collected and mitochondrial fractions were isolated using a previously reported protocol (*42*). Briefly, cell membranes were physically broken with a douncer in 500 μl detergent-free buffer A (20 mM HEPES, 220 mM Mannitol, 70 mM Sucrose, 1 mM EDTA, 2 mg/ml BSA) and lysates were centrifuged at 200 x g for 5 min at 4 °C in a swing-rotor. Supernatants were collected (total lysate) and centrifuged at 12,000 x g for 10 min at 4 °C. Supernatants were discarded and the pellet containing the crude mitochondrial fraction was resuspended in 100 µl buffer B (20 mM HEPES, 220 mM Mannitol, 70 mM Sucrose, 1 mM EDTA) and centrifuged at 12,000 x g for 15 min at 4 °C. Pellets were resuspended in either 100 µl buffer B alone or containing 3 µl thermosensitive Proteinase K (PK; NEB) and incubated at 4 °C for 30 min. PK was inactivated at 65 °C for 15 min. Supernatants were mixed with Laemmli sample buffer and incubated at 65 °C for 10 min. Samples were electrophoretically separated by SDS-PAGE (10 % polyacrylamide gel) and immunoblotted on Nitrocellulose membranes. Antibodies employed for immunoblotting included anti-GCN4 rabbit antibody (1:10,000; Novus, #NBP2-81274) and anti-ATP5α mouse antibody (1:1000; Invitrogen, #43-9800).

In all cases, secondary antibodies anti-mouse or anti-rabbit Fc chains generated in either goat or donkey were fluorescently conjugated with either A488 or A657, and detected with an iBright imaging system (Thermo Fisher Scientific).

## Live cell mRNA imaging

Live cell imaging of *Pink1* mRNA in mouse primary hippocampal neurons was performed as described previously (*13,14*). Neurons were seeded in 24-well glass bottom plates (CellVis) coated with PLL and laminin at a density of 1-1.5x10^5^ cells per well and cultured in NB+B27+PSG as described above. On DIV7, transfection was performed with Lipofectamine® 2000 transfection reagent (Thermo Fisher Scientific) introducing 4 µg of Pink1-kinase dead-MS2-PP7 and 1 µg of nls-ha-MCP-VenusN-nls-ha-PCP-VenusC along with 0.3 µg of TMEM192-mRFP-3xHA into the neurons. On DIV9, the neuronal medium was exchanged with Hibernate E medium without phenol red (BrainBits) and the neurons were incubated with 20 µM Compound C (CC; Abcam, 2 h) or 500 nM insulin (Sigma-Aldrich, 1 h). Afterwards, the neurons were imaged at the Imaging Facility of the Max Planck Institute for Biological Intelligence, Martinsried, Germany, using an Eclipse Ti2 spinning disk microscope (Nikon) with a DS-Qi2 high-sensitivity monochrome camera, a 60×/NA 1.2 oil immersion objective as well as the NIS-Elements software (version 5.21.03).

## Proximity Ligation Assay

Mouse primary hippocampal neurons were seeded on acid-washed glass coverslips (1.5 mm, Marienfeld) coated with PLL and laminin at a density of 5x10^4^ cells per well and cultured in NB+B27+PSG as described above. For detecting a PLA signal between the PINK1 SunTag signal (scFv-sfGFP) and the ribosomes, RRBP1 or Sec61b, on DIV5 primary neurons were transfected with 0.25 µg of scFv-sfGFP, with or without 1 µg Pink1-SunTagcyto, and with or without 0.3 µg control shRNA or DNAJB6 shRNA (depending on the experiment) using Lipofectamine® 2000 transfection reagent (Thermo Fisher Scientific). On DIV7, the neurons were incubated with or without 20 µM CC (Abcam) for 2 h and with or without 20 µM Antimycin A (AA, Sigma-Aldrich; 45 min), followed by fixation using 4 % warm PFA for 15 min at room temperature. The neurons were permeabilized in 0.3 % Triton X-100/PBS for 10 min and the next steps were conducted following the manufacturer’s guidelines (Sigma-Aldrich) and as previously described (*13,14*). Neurons were first incubated with the Duolink blocking solution for 60 min at 37 °C in a humidity chamber and afterwards with the primary antibody anti-RPS6 rabbit (5G10) (1:200; Cell Signaling Technology, #2217), anti-RRBP1 rabbit (1:500; Abcam, #ab95983) or anti-Sec61b rabbit (1:500; Proteintech, #15087-1-AP) diluted in Duolink antibody diluent at 4 °C overnight. The expressed scFv-sfGFP serves as the second primary antibody (mouse) inside the cells. One of the control samples was incubated without the anti-RPS6 antibody. On the following day, cells underwent two 5-min washes at room temperature with buffer A (0.01 M Tris, 0.15 M NaCl, 0.05 % Tween 20) followed by incubation with the Duolink anti-rabbit minus and anti-mouse plus PLA probes, diluted in a 1:5 ratio in the Duolink antibody diluent, for 60 min at 37 °C. After another two 5-min washes with buffer A at room temperature, a ligase, diluted in a 1:40 ratio in the Duolink ligation buffer, was added to the cells for 30 min at 37 °C. Afterwards, cells were washed twice for 5 min with buffer A at room temperature and subsequently treated with a polymerase, diluted in a 1:80 ratio in the Duolink amplification buffer containing fluorescently-labeled oligonucleotides (Cy5) for 100 min at 37 °C. After two 10-min washing steps in buffer B (0.2 M Tris, 0.1 M NaCl) and a final 1-min washing step in 0.01x buffer B at room temperature, Fluoromount G (Invitrogen) was used to mount the coverslips. The coverslips were imaged at the Imaging Facility of the Max Planck Institute for Biological Intelligence, Martinsried, Germany, using an Eclipse Ti2 spinning disk microscope (Nikon) with a DS-Qi2 high-sensitivity monochrome camera, a 60×/NA 1.2 oil immersion objective as well as the NIS-Elements software (version 5.21.03).

## Protein translation assays

### SunTag system

Mouse primary hippocampal neurons were seeded in 24-well glass bottom plates (CellVis) coated with PLL and laminin at a density of 1-1.5x10^5^ cells per well. HeLa cells were seeded in 24-well glass bottom plates (CellVis) at a density of 5x10^4^ cells per well. Cells were cultured in their respective media, as described above. On DIV7 or DIV9, neuronal transfection was performed with Lipofectamine® 2000 transfection reagent (Thermo Fisher Scientific) introducing 1 µg Pink1-SunTagmito or Pink1-SunTagcyto and 0.25 µg of scFv-sfGFP along with 0.3 µg of the respective organellar markers (mito-BFP, mito-mRaspberry-7, mCherry-Rab7A, TMEM192- mRFP-3xHA or pLAMP1-mCherry) and, when indicated, 0.3 µg of plasmids encoding mycSYNJ2BP-WT or -S21E, control shRNA, DNAJB6 shRNA or RRBP1 shRNA into the neurons. One day after seeding, HeLa cells were transfected with 1 µg Pink1-SunTagcyto and 0.25 µg of scFv-sfGFP using calcium phosphate. On DIV9-DIV12 or two days after transfection, the neuronal and HeLa cell medium, respectively, was exchanged with Hibernate E medium without phenol red (BrainBits) and the cells underwent the following treatments: 20 µM CC (Abcam, 2 h), 1 mM AICAR (Abcam; 2 h), 500 nM insulin (Sigma-Aldrich; 1 h), 1 µM Wortmannin (EMD Millipore; 2 h), 20 µM Antimycin A (AA, Sigma-Aldrich; 45 min), or 100- 200 µg/ml Puromycin (Enzo Life Sciences; 30 min-2 h). Afterwards, the neurons and HeLa cells were imaged in live-time at the Imaging Facility of the Max Planck Institute for Biological Intelligence, Martinsried, Germany, using an Eclipse Ti2 spinning disk microscope (Nikon) with a DS-Qi2 high-sensitivity monochrome camera, a 60×/NA 1.2 oil immersion objective as well as the NIS-Elements software (version 5.21.03). Cells were kept at a controlled temperature of 37 °C during the imaging session. For time-lapse imaging, cells were recorded every minute, for a total of 45 minutes. To image the same cells before and after the indicated treatments, the cellular position was stored using the NIS-Elements software (Nikon).

## Proximity Ligation Assay coupled with puromycylation (puro-PLA)

Human iPSC-derived neurons were seeded on acid-washed glass coverslips (1.5 mm, Marienfeld) coated with PLL and laminin at a density of 5x10^4^ cells per well and cultured in conditioned Südhof neuronal growth medium as described above. On DIV14, the following treatments were performed: 20 µM CC (Abcam, 2 h), 1 mM AICAR (Abcam, 2 h), 500 nM insulin (Sigma-Aldrich, 1 h) or conditioned Südhof neuronal growth medium lacking insulin (2 h). Afterwards, 10 µg/ml Puromycin (Enzo Life Sciences) was added to the medium of the cells for 5 min at 37 °C. To stop the puromycylation reaction, cells were briefly washed with PBS containing 1 mM MgCl2, 0.1 mM CaCl2, and 10 µg/ml cycloheximide (Enzo Life Sciences) followed by fixation using 4 % warm PFA for 15 min at room temperature. The proximity ligation assay was performed as described above including permeabilization, blocking, primary antibody incubation, PLA probe incubation, ligase treatment, and polymerase treatment. The following primary antibodies were used: anti-PINK1 rabbit antibody (1:1,000, Novus Biologicals, #BC100-494), anti-Puromycin mouse (3RH11) antibody (1:1,000, Kerafast, #EQ0001). After mounting of the coverslips using Fluoromount G (Invitrogen), they were imaged at the Imaging Facility of the Max Planck Institute for Biological Intelligence, Martinsried, Germany, using an Eclipse Ti2 spinning disk microscope (Nikon) with a DS-Qi2 high-sensitivity monochrome camera, a 60×/NA 1.2 oil immersion objective as well as the NIS-Elements software (version 5.21.03).

## BirA assay

Mouse primary cortical neurons were seeded in 6-well plates (Greiner) coated with PLL and laminin at a density of 2x10^6^ cells per well. On DIV1, neurons were lentivirally transduced for 5- 8 h with the AviTag-RPL10a and the rtTA as well as one of the following BirA constructs: BirA-mCherry-Fis1, BirA-mCherry-Sec61β, BirA-mCherry-Rab7 or cytosolic BirA-mCherry. On DIV5, neurons were treated with 2 ng/µl doxycycline for 4 h to induce expression of the BirA constructs. Additionally, 20 µM CC (Abcam, 2 h) was added to the neurons. Subsequently, neurons were lysed using a buffer containing 25 mM Tris/HCl pH 7.4, 1 % NP-40, 150 mM NaCl, supplemented with 1X protease inhibitor cocktail (Roche), 200 µM PMSF, and 0.1% RNasin (Promega). After centrifugation at 9,300 x g for 1 min, the pellets were discarded and 10 % of the supernatant was kept as input, while the remaining cell lysate was incubated with 30 µl Streptavidin-Agarose beads (Sigma-Aldrich) per sample for 40 h at 4 °C. Afterwards, the beads were washed five times in columns (MoBiTec) in a buffer composed of 20 mM Tris/HCl pH 8, 5 mM MgCl2, 140 mM NaCl, 0.1 % Triton X-100, 200 µM PMSF and 0.1% RNasin. The elution was performed by adding Laemmli sample buffer followed by boiling at 95 °C for 2 min.

For RNA isolation, input as well as pulldown samples were mixed with 1 ml TRIzol (Sigma-Aldrich) followed by addition of 100 µl BCP (1-Bromo-3-chloropropane) per sample. After centrifugation for 15 min at 12,000 x g at 4 °C, the upper aqueous phase was transferred to a new tube containing one volume BCP. After a second centrifugation for 5 min at 12,000 x g at 4 °C, the upper phase was mixed with 2.5 volumes of 100 % ethanol, 100 µl 3 M sodium acetate and 10 µl glycogen. The samples were incubated at -20 °C for at least 30 min and then centrifuged for 20 min at 12,000 x g at 4 °C. The supernatant was discarded and the pellet was resuspended in 250 µl MilliQ water followed by addition of 1 ml 100 % ethanol, 10 µl glycogen and 100 µl 3 M sodium acetate. After a final centrifugation of 5 min at 7,500 x g at 4 °C, the pellet was resuspended in 20 µl RNAse-free water.

The RNA samples were analyzed by RT-qPCR to determine *Pink1* transcript levels. 200 ng were used for complementary DNA (cDNA) synthesis with qScript^TM^ cDNA SuperMix (Quantabio). The RT-qPCR assay was conducted in a Mic (magnetic induction cycler) PCR machine (Bio Molecular Systems) utilizing the PerfeCTa SYBR® Green FastMix (Quantabio). The specific primers used for the assay were as follows (5’ → 3’): Pink1 forward: AATGAGCCAGGAGCTGGTC; Pink1 reverse: GTACTGGCGCAGGGTACAG; β-actin forward: ACACTGTGCCCATCTACG; β-actin reverse: GCTGTGGTGGTGAAGCTGTAG.

Quantification of the mRNA enrichment at the indicated subcellular localizations and treatments was evaluated following the percent input method of the Ct values.

## Mitophagy assays

### Optineurin and phospho-ubiquitin immunostaining

Mouse primary hippocampal neurons were seeded on acid-washed glass coverslips (1.5 mm, Marienfeld) coated with PLL and laminin at a density of 5x10^4^ cells per well and cultured in NB+B27+PSG as described above. On DIV5, transfection was performed with Lipofectamine® 2000 transfection reagent (Thermo Fisher Scientific) introducing 0.3 µg mito-meGFP as well as 0.3 µg control shRNA, RRBP1 shRNA or DNAJB6 shRNA into the neurons. On DIV8, neurons were incubated for 45 min with or without 20 µM AA (Sigma-Aldrich) followed by fixation using 4 % warm PFA for 15 min at room temperature. The neurons were permeabilized in 0.3 % Triton X-100/PBS for 10 min, blocked in 1 % BSA/PBS for 1 h at room temperature, and finally incubated with the primary antibodies (anti-optineurin rabbit antibody (1:500, Abcam, #ab23666); anti-phospho-ubiquitin rabbit antibody (1:200, Millipore, #ABS1513-I); anti-βIII tubulin mouse (2G10) antibody (1:1,000; Invitrogen, #MA1-118)) diluted in 1 % BSA/PBS overnight at 4 °C. On the following day, the fixed neurons were subjected to three 5-min washes in PBS at room temperature and then incubated with the secondary fluorescent antibodies Alexa Fluor 568 goat anti-mouse and Alexa Fluor 647 goat anti-rabbit (Invitrogen, 1:500) diluted in 1 % BSA/PBS for 2 h at room temperature. After additional three 5-min washes in PBS, the coverslips were mounted using Fluoromount G (Invitrogen) and imaged at the Imaging Facility of the Max Planck Institute for Biological Intelligence, Martinsried, Germany, using an Eclipse Ti2 spinning disk microscope (Nikon) with a DS-Qi2 high-sensitivity monochrome camera, a 60×/NA 1.2 oil immersion objective as well as the NIS-Elements software (version 5.21.03).

## Mitochondrial and lysosomal colocalization assay

Mouse primary hippocampal neurons were seeded in 24-well glass bottom plates (CellVis) coated with PLL and laminin at a density of 10^5^ cells per well and cultured in NB+B27+PSG as described above. On DIV6, transfection was performed with Lipofectamine® 2000 transfection reagent (Thermo Fisher Scientific) introducing 0.3 µg mito-meGFP and 0.3 µg LAMP1-mCherry as well as 0.3 µg control shRNA or DNAJB6 shRNA into the neurons. On DIV9, the neuronal medium was exchanged with Hibernate E medium without phenol red (BrainBits) and the neurons were incubated for 45 min with or without 20 µM AA (Sigma-Aldrich). Afterwards, the neurons were live-time imaged at the Imaging Facility of the Max Planck Institute for Biological Intelligence, Martinsried, Germany, using an Eclipse Ti2 spinning disk microscope (Nikon) with a DS-Qi2 high-sensitivity monochrome camera, a 60×/NA 1.2 oil immersion objective as well as the NIS-Elements software (version 5.21.03).

## DNAJB6 immunostaining

Mouse primary hippocampal neurons were seeded on acid-washed glass coverslips (1.5 mm, Marienfeld) coated with PLL and laminin at a density of 5x10^4^ cells per well. HeLa cells were seeded on acid-washed glass coverslips (1.5 mm, Marienfeld) at a density of 5x10^4^ cells per well. Primary neurons and HeLa cells were cultured in NB+B27+PSG and DMEM containing 10 % FBS and 1 % P/S, respectively, as described above. On DIV6, transfection of primary neurons was performed with Lipofectamine® 2000 transfection reagent (Thermo Fisher Scientific) introducing 1 µg of Pink1-SunTagcyto and 0.25 µg of scFv-sfGFP into the cells. On DIV8 or two days after seeding, neurons and HeLa cells, respectively, were incubated with or without 20 µM CC (Abcam) for 2 h followed by fixation using 4 % PFA for 15 min at room temperature. The cells were permeabilized in 0.3 % Triton X-100/PBS for 10 min, blocked in 1 % BSA/PBS for 1 h at room temperature, and finally incubated with the primary antibodies (anti-DNAJB6 rabbit antibody (1:500, Proteintech, #11707-1-AP); anti-βIII tubulin mouse (2G10) antibody (1:1,000, Invitrogen, #MA1-118); anti-β-actin mouse (AC-74) antibody (1:1,000, Sigma-Aldrich, #A5316)) diluted in 1 % BSA/PBS overnight at 4 °C. On the following day, the fixed cells were subjected to three 5- min washes in PBS at room temperature and then incubated with the secondary fluorescent antibodies Alexa Fluor 647 goat anti-rabbit and Alexa Fluor 488 goat anti-mouse (Invitrogen, 1:500) diluted in 1 % BSA/PBS for 2 h at room temperature. After additional three 5-min washes in PBS, the coverslips were mounted using Fluoromount G (Invitrogen) and imaged at the Imaging Facility of the Max Planck Institute for Biological Intelligence, Martinsried, Germany, using an Eclipse Ti2 spinning disk microscope (Nikon) with a DS-Qi2 high-sensitivity monochrome camera, a 60×/NA 1.2 oil immersion objective as well as the NIS-Elements software (version 5.21.03).

## Correlative light and scanning electron microscopy

For correlative imaging experiments, mouse primary hippocampal neurons were seeded on patterned 35 mm glass bottom dishes (MatTek, 14 mm glass diameter, 1.5 gridded coverslip) coated with PLL and laminin at a density of 1.5x10^5^ cells per dish and cultured as described above. On DIV7, transfection was performed with Lipofectamine® 2000 transfection reagent (Thermo Fisher Scientific) introducing 1 µg of Pink1-SunTagcyto, 0.25 µg of scFv-sfGFP, 0.3 µg of mito-BFP, and 0.3 µg of TMEM192-mRFP-3xHA into the neurons. On DIV9, imaging was performed at the Imaging Facility of the Max Planck Institute for Biological Intelligence, Martinsried, Germany, using an Eclipse Ti2 spinning disk microscope (Nikon) with a DS-Qi2 high-sensitivity monochrome camera, a 60×/NA 1.2 oil immersion objective as well as the NIS-Elements software (version 5.21.03). Before imaging, the neuronal medium was exchanged with Hibernate E medium without phenol red (BrainBits) and the neurons underwent a 2 h treatment with 20 µM CC (Abcam). Following imaging, immediate fixation was performed using a freshly prepared 2.5 % glutaraldehyde solution in 0.1 M cacodylate buffer (Science Services). The 2x fixative solution, pre-warmed to 37 °C, was added directly into the Hibernate E medium of the neurons at a 1:1 ratio.

Following a 5 min incubation at 37 °C, it was replaced with 1x fixative and further incubated for 25 min on ice. Removal of the fixative involved three 5 min washes with 0.1 M cacodylate buffer on ice. Further processing was performed at the electron microscopy hub at the German Center for Neurodegenerative Diseases (DZNE), Munich, Germany. Employing a reduced osmium thiocarbohydrazide osmium (rOTO) *en bloc* staining protocol, the samples underwent a sequential series of treatments. This process involved postfixation in a solution containing 2 % osmium tetroxide (Science Services), 1.5 % potassium ferricyanide (Sigma-Aldrich) in 0.1 M sodium cacodylate buffer (pH 7.4; Science Services) (*43*). The staining was intensified through a 45 min incubation at 40 °C with 1 % thiocarbohydrazide (Sigma-Aldrich). Subsequently, the neurons underwent rinsing in water and immersion in a 2 % aqueous solution of osmium tetroxide. After another wash, an additional level of contrast was achieved through an overnight incubation in a 1 % aqueous solution of uranyl acetate at 4 °C, followed by an extra 2 h incubation at 50 °C. In preparation for subsequent processing steps, the samples underwent dehydration through a series of increasing ethanol concentrations and were infiltrated with LX112 (LADD). Following the curing process, the block was removed from the glass coverslip through a series of freeze-thaw cycles. The region of interest was trimmed (TRIM2, Leica) in accordance with the previously acquired grid pattern from confocal imaging. Serial sections, with a nominal thickness of 80 nm, were acquired using a 35° ultra-diamond knife (Diatome) on an ATUMtome (Powertome, RMC). These sections were collected onto carbon-coated Kapton tape (kindly provided by Jeff Lichtman and Richard Schalek), freshly treated with plasma (custom-built, based on PELCO easiGlow, adopted from M. Terasaki, U. Connecticut, CT). The tape stripes were affixed onto adhesive carbon tape (Science Services), and this assembly was then secured onto 4-inch silicon wafers (Siegert Wafer), and grounded through adhesive carbon tape stripes (Science Services). EM micrographs were acquired using an Apreo S2 SEM (Thermo Fisher Scientific) as previously described (*44*). Hierarchical imaging of serial sections was achieved through initial mapping of the plastic tape stripes at low lateral resolution and capture of the entire tissue sections at a medium resolution (100-200 nm). Correlation of the region of interest was established through the integration of the grid pattern and neuronal morphology. Subsequently, micrographs of serial sections were acquired at a lateral resolution of 5 nm. To align the serial section imaging data, a hybrid approach combining both automated and manual processing procedures was employed within Fiji TrakEM2 (44). Following the alignment, single sections containing the organellar structures of interest were chosen.

## QUANTIFICATION AND STATISTICAL ANALYSIS

The presented data are expressed as mean ± SEM, unless otherwise stated. Graphical representation and statistical analysis were conducted using GraphPad Prism (version 9.1.0) for Windows (GraphPad Software, San Diego, California, USA, www.graphpad.com) or MS Excel. To compare two conditions a Student’s t-test was employed, while a one-way ANOVA with Tukey’s multiple comparisons was done for comparison of multiple groups and a two-way ANOVA with Tukey’s or Sidak’s multiple comparisons was used for comparison of multiple paired groups. The statistical significance thresholds were set as follows: p<0.05 (*), p<0.01 (**), p<0.001 (***) and p<0.0001 (****). For statistically non-significant comparisons, the p-values are explicitly indicated in the figures.

Immunoblot images were obtained using the Invitrogen iBright FL1000 Imaging System (Thermo Fisher Scientific) and the intensity of the bands was analyzed in the Fiji distribution of ImageJ (National Institutes of Health) (*45*). Microscopy data quantification was carried out using Fiji/ImageJ. Researchers were blind to the experimental conditions. Images are displayed in grayscale (with signal inversion) when assessing the appearance of specific structures (SunTag clusters, PLA dots) involving a single channel. However, when the analysis involved colocalization between different signals/channels, the images were presented in color. For analysis of number of clusters, intensity and area of clusters, maximum z-projection was performed on a square-shaped area including the entire cell body. Images were thresholded for the GFP channel (SunTag signal). For the analysis of the number of clusters, the function ‘find edges’ in Fiji/ImageJ was used to define the contour of the clusters. The identification of the SunTag clusters was performed using the particle analyzer in Fiji/ImageJ. Subsequently, the intensity as well as the area of the non-thresholded images were measured. For analysis of colocalization, the JaCoP plugin (*46*) in Fiji/ImageJ was used to determine the Manders’ colocalization coefficient between the GFP (SunTag signal) and the organelle channel (mitochondria, endolysosomes) in z-stack images containing the entire cell body without maximum z-projection. For the rotated GFP (SunTag signal) quantification, a square (10 x 10 µm) within the cell body was selected based on the organelle channel and excluding the nucleus. The Manders’ colocalization coefficient was determined both before and after rotating the GFP channel 90°. The Manders’ coefficient values, mean gray values (intensities) and area values were exported to Excel and plotted using GraphPad Prism. To quantify *Pink1* mRNA imaging, the Manders’ colocalization coefficient between the Venus and the endolysosomal channel was determined in z-stack images using the JaCoP plugin in Fiji/ImageJ as described before (*13,14*). A square-shaped area including the entire cell body was chosen. The resulting Manders’ colocalization coefficients were exported to Excel and plotted using GraphPad Prism. To analyze the optineurin and phospho-ubiquitin stainings, the Manders’ colocalization coefficient between the phospho-ubiquitin/optineurin and the mitochondrial channel was determined using the JaCoP plugin in Fiji/ImageJ as described before.^19,20^ Images of intact neurites, identified by the βIII-tubulin signal, were straightened with a 20-pixel or 50-pixel margin following maximum z-projection. The resulting Manders’ colocalization coefficients were exported to Excel and plotted using GraphPad Prism.

## Acknowledgments

We thank C. Weiß, J. Lindner, M. Martins, C. Niemann and G. Kislinger for technical support and the members of the Harbauer laboratory for their support and many fruitful discussions. We are grateful to R. Kasper/E. Laurell/C. Polisseni and M. Spitaler/M. Oster from the Imaging facilities of the MPIs for Biological Intelligence and Biochemistry, respectively, for assistance with live cell imaging.

## Funding

Work in **ABH’s** lab is supported by Max Planck Society

DFG (DFG, HA 7728/2-1 - ID 453679203, TRR353 – ID 471011418)

European Union (ERC StG Project 101077138 — MitoPIP)

Germany’s Excellence Strategy within the framework of the Munich Cluster for Systems Neurology (EXC 2145 SyNergy – ID 390857198).

Work in **TM’s** lab is supported by German Research Foundation (DFG, Mi 694/8-1 “, FOR Immunostroke (Mi 694/9-1/2 – ID 428663564, TRR 274/1 Projects C02, Z01, respectively – ID 408885537).

German Center for Neurodegenerative Diseases (DZNE).

DFG instrumentation grant (INST95/1755-1 FUGG, ID 518284373).

Chan Zuckerberg Initiative (CZI) grant for Creating Community & Building Capacity.

Germany’s Excellence Strategy within the framework of the Munich Cluster for Systems Neurology (EXC 2145 SyNergy – ID 390857198).

## Author Contributions

Conceptualization: ABH Methodology: ABH, JTH, IS, AS, MS Investigation: JTH, IS, AS, MS Visualization: JTH, IS, TM Supervision: ABH, TM Writing – original draft: ABH, JTH, IS Writing – review & editing: all authors

## Declaration of Interests

The authors declare no competing interests.

**Data and materials availability:** Requests for resources and reagents should be directed to and will be fulfilled by the lead contact Angelika B. Harbauer.

Microscopy data reported in this paper will be shared by the lead contact upon request. Any additional information required to reanalyze the data reported in this paper is available from the lead contact upon request.

**Supplementary Materials** Materials and Methods Supplementary Text

Figs. S1 to S# Tables S1 to S# References (*##*–*##*) Movies S1 to S# Audio S1 to S# Data S1 to S#

---

**Fig. S1.**
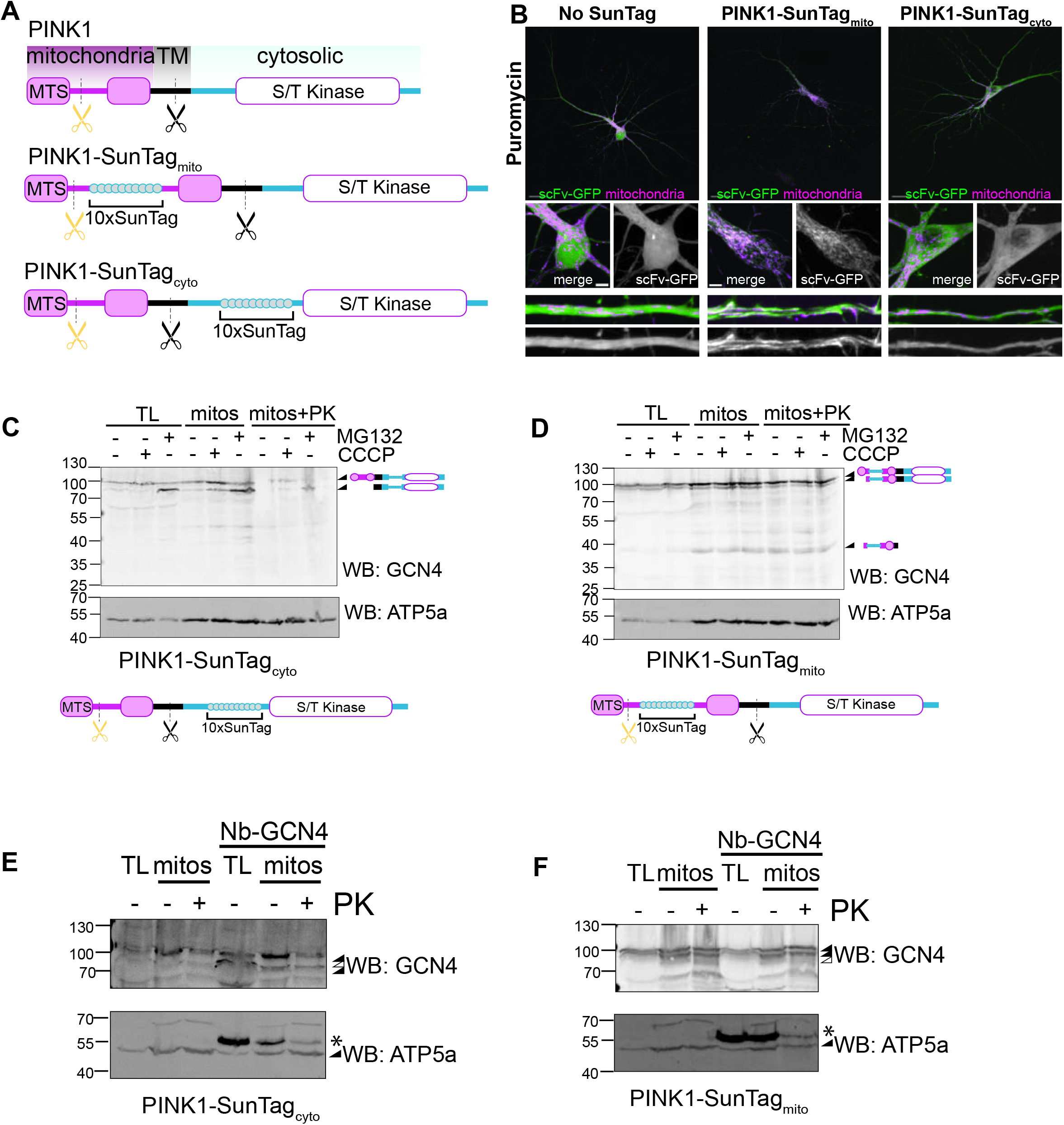
PINK1 localization visualized by the SunTag system. **A** Schematic representation of PINK1 protein domains and the PINK1-SunTagmito and PINK1- SunTagcyto reporters. A yellow scissor indicates MPP cleavage, a black scissor indicates a cleavage within the transmembrane segment by PARL. MTS=mitochondrial targeting sequence; TM=transmembrane domain; S/T=Serine/Threonine. **B** Representative images showing the distribution of either the nanobody scFv-GFP alone or cotransfected with either PINK1-SunTagmito or PINK1-SunTagcyto reporters in mouse hippocampal neurons (green). Mitochondria are visualized by coexpression with Cox8-BFP protein (magenta). Puromycin treatment was performed for 30 min at a concentration of 100 µg/ml. Inserts showing magnified somas and dendrites are included below. Black-white images correspond to the scFv-GFP signal only. **C,D** Western blots of total lysates (TL) and mitochondrial fractions (mitos) obtained from HEK293 cells expressing the PINK1-SunTagcyto (**C**) or PINK1-SunTagmito (**D**) reporters after the indicated treatments (100 µM CCCP or 10 µM MG132 for 2 h). Reporters were detected with an anti-GCN4 antibody directed against the SunTag epitopes. Isolated mitochondria were treated with proteinase K (PK) when indicated. Arrowheads point at the bands corresponding to full length and processed PINK1-SunTagcyto (**C**) or PINK1-SunTagmito (**D**) with the corresponding schematic representation. The anti-ATP5a signal was used as loading control of the mitochondrial content. **E,F** Western blots of total lysates (TL) and mitochondrial fractions (mitos) obtained from HEK293 cells expressing the PINK1-SunTagcyto (**E**) or PINK1-SunTagmito (**F**) reporters alone or together with intracellular expression of the nanobody scFv-GFP (Nb-GCN4). Reporters were detected with an anti-GCN4 antibody directed against the SunTag epitopes. Isolated mitochondria were treated with proteinase K (PK) when indicated. The anti-ATP5a signal was used as loading control of the mitochondrial content. Black arrowheads point at the bands corresponding to full length and processed PINK1-SunTag (upper panels) or ATP5a (lower panels). White arrowheads indicate the expected size of processed PINK1-SunTag that was not observed. Asterisks denote the co-expressed nanobody, detected by the anti-mouse secondary antibody. Scale bars, 10 µm for whole soma, 2 µm for insets.

**Fig. S2.**
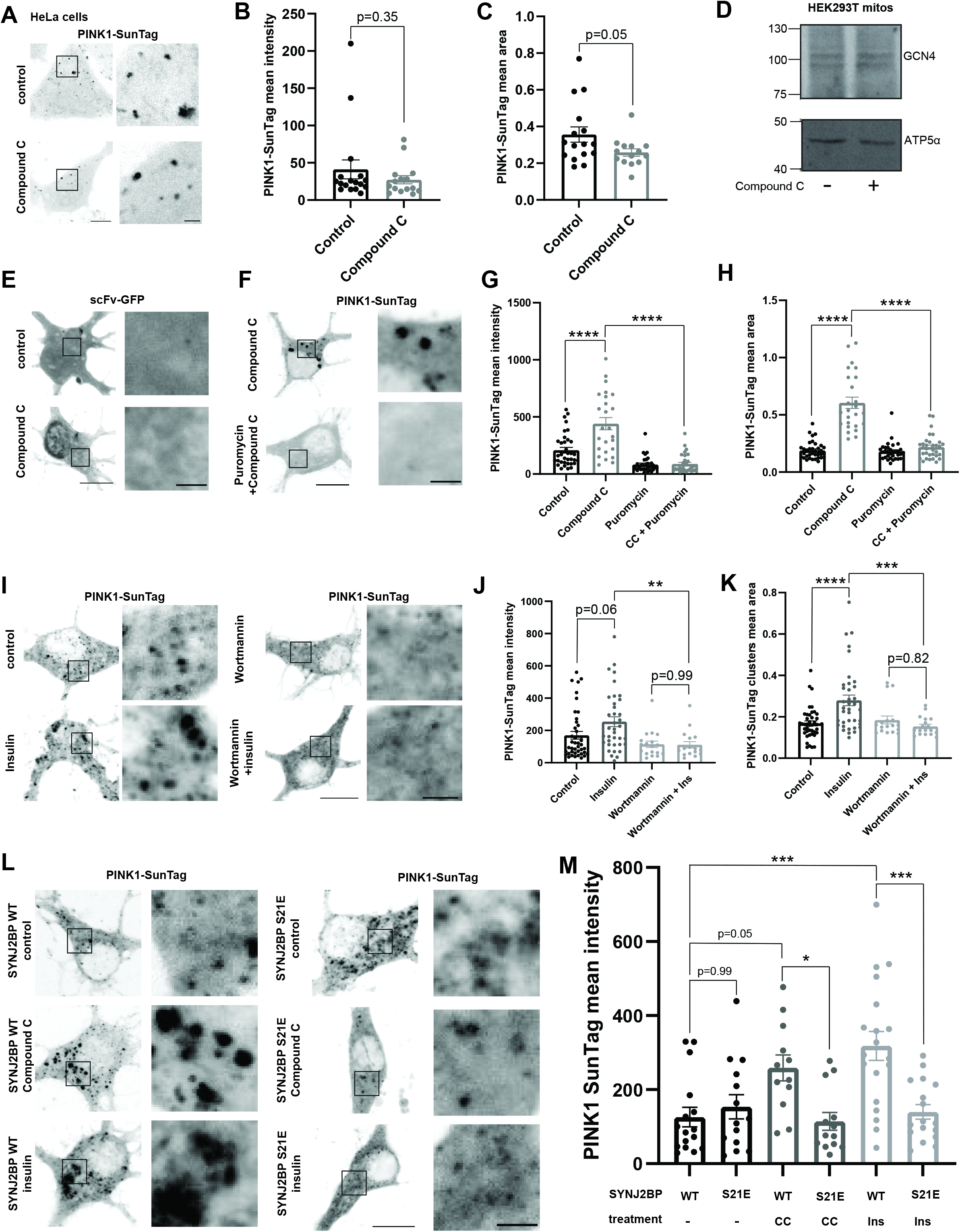
Cytosolic PINK1-SunTag cluster formation also occurs upon insulin treatment and is neuron-specific. **A** Representative images of cytosolic PINK1 clusters visualized by the SunTag system upon control (untreated) or CC (20 µM, 2 h) treatment in HeLa cells. **B** Quantification of the PINK1- SunTag protein cluster mean intensity as in **A**. Student’s t-test; n = 15-17. **C** Quantification of the PINK1 protein cluster mean area as in **A**. Student’s t-test; n = 14-16. **D** Representative immunoblot image of isolated mitochondria from HEK293T cells overexpressing the PINK1-SunTag construct following control (untreated) or CC (20 µM, 2 h) treatment. The reporter was detected with an anti-GCN4 antibody directed against the SunTag epitopes. The anti-ATP5α signal was used as loading control of the mitochondrial content. **E** Representative images of hippocampal neurons overexpressing scFv-GFP alone (without the plasmid encoding PINK1-SunTag) upon control (untreated) or CC (20 µM, 2 h) treatment. **F** Representative images of cytosolic PINK1 protein clusters visualized by the SunTag system upon CC (20 µM, 2 h) or simultaneous treatment with CC and Puromycin (200 µg/ml, 2 h) in hippocampal neurons. **G** Quantification of the PINK1 protein cluster mean intensity following Puromycin and CC treatment as in **F**. One-way ANOVA followed by Tukey’s post hoc test; n = 25-35; p < 0.0001 (****). **H** Quantification of the PINK1 protein cluster mean area following Puromycin and CC treatment as in **F**. One-way ANOVA followed by Tukey’s post hoc test; n = 25-35; p < 0.0001 (****). **I** Representative images of cytosolic PINK1 clusters visualized by the SunTag system upon insulin (500 nM, 1 h) treatment, with vehicle (control) or with Wortmannin (phosphatidylinositol 3-kinase inhibitor, 1 µM, 2 h) pre-treatment in hippocampal neurons. **J** Quantification of the PINK1 protein cluster mean intensity in hippocampal neurons treated as in **I**. One-way ANOVA followed by Tukey’s post hoc test; n = 18-42, p < 0.01 (**). **K** Quantification of the PINK1 protein cluster mean area in hippocampal neurons treated with insulin (500 nM, 1 h) and pre-treated with Wortmannin (1 µM, 2 h) as in **I**. One-way ANOVA followed by Tukey’s post hoc test; n = 18-42, p < 0.001 (***), p < 0.0001 (****). **L** Representative images of cytosolic PINK1 protein clusters visualized by the SunTag system in hippocampal neurons overexpressing either SYNJ2BP WT or S21E and treated with either control (untreated), CC (20 µM, 2 h) or insulin (ins, 500 nM, 1 h). **M** Quantification of the PINK1 protein cluster mean intensity as in **L**. One-way ANOVA followed by Tukey’s post hoc test; n = 12-19; p < 0.05 (*), p < 0.001 (***). Data are expressed as mean±SEM. All data points correspond to single cells coming from ≥3 biological replicates (**B,C,G,H,J,K,M**). Scale bars, 10 µm for whole soma, 2 µm for insets.

**Fig. S3.**
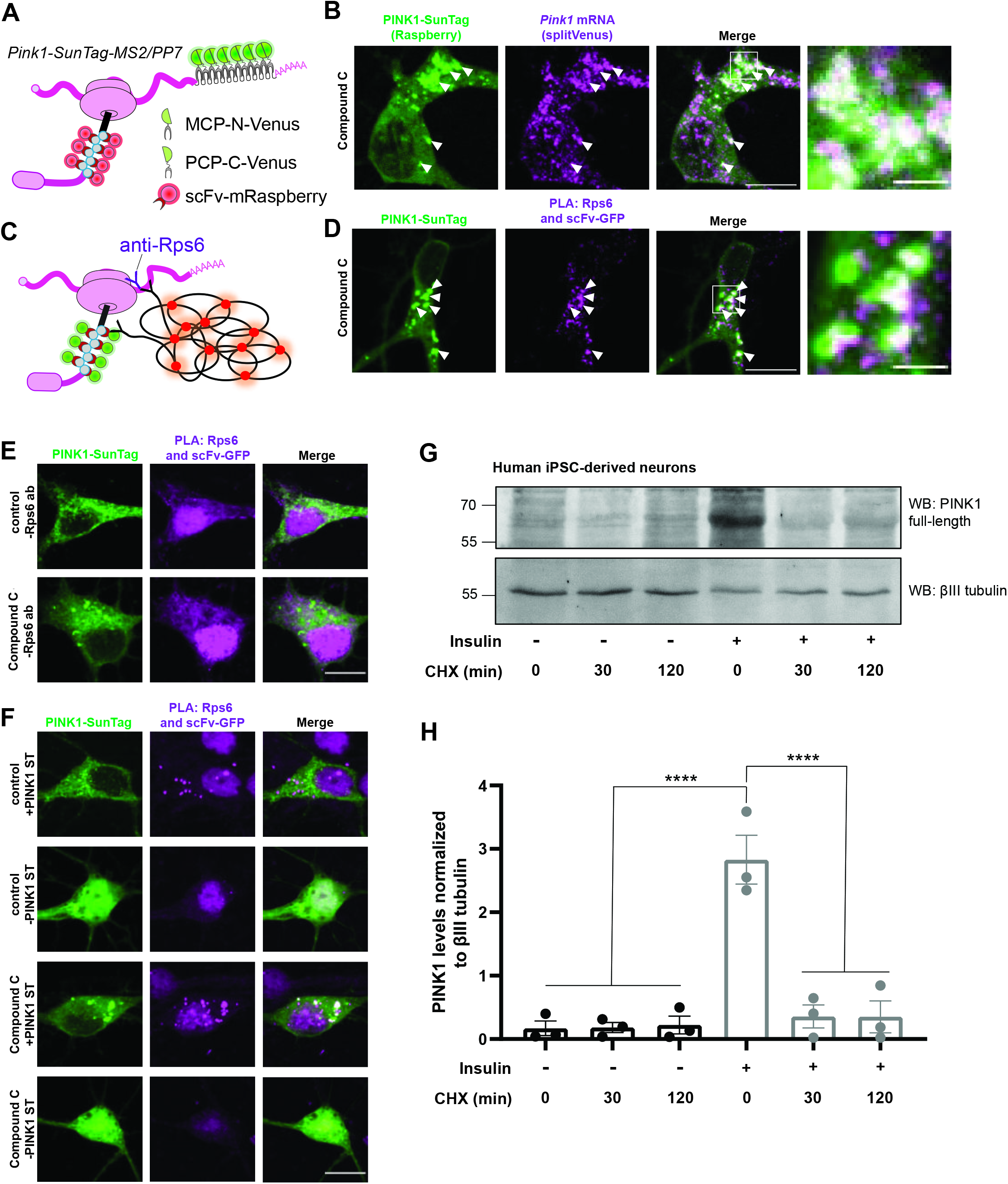
*Pink1* mRNA and ribosomes localize to PINK1-SunTag clusters and PINK1 degradation is not impaired upon insulin. **A** Schematic illustrating the dual imaging of the *Pink1* mRNA (MS2/PP7; green) and the newly synthesized PINK1 protein (SunTag; red). **B** Representative images of cytosolic PINK1 clusters visualized by the SunTag system (using scFv-mRaspberry instead of scFv-GFP, pseudocolored in green for consistency) and *Pink1* mRNA visualized by the MS2/PP7-split Venus approach (magenta) upon CC (20 µM, 2 h) treatment in hippocampal neurons. Arrowheads point to regions where the *Pink1* mRNA and the PINK1-SunTag clusters overlap. **C** Schematic illustrating the proximity ligation assay for visualizing cytosolic PINK1-SunTag clusters in close proximity to Rps6-positive ribosomes. **D** Representative images of cytosolic PINK1 clusters visualized by the SunTag system as well as the PLA signal for scFv-GFP and Rps6 upon CC (20 µM, 2 h) treatment in hippocampal neurons. Arrowheads point to regions where the PINK1-SunTag clusters and the PLA signal overlap. **E** Representative images of cytosolic PINK1 clusters visualized by the SunTag system upon control (untreated) or CC (20 µM, 2 h) treatment in hippocampal neurons. The PLA signal is shown for scFv-GFP but without addition of the anti-Rps6 antibody (-Rps6 ab) and acts as a control for Figure **S3D**. Note that there is no PLA signal in this condition. **F** Representative images of hippocampal neurons overexpressing scFv-GFP with (+) or without (-) PINK1-SunTag reporter (PINK1-ST) upon control (untreated) or CC (20 µM, 2 h) treatment. The PLA signal is shown for scFv-GFP and Rps6 and acts as a control for Figure **S3D**. Note that there is no PLA signal without expression of PINK1-SunTag reporter, despite the diffuse localization of the scFv-GFP nanobody. **G** Representative immunoblot image of lysates from human iPSC-derived neurons cultured in the absence or presence of insulin for 2 h followed by treatment with cycloheximide (70 µM) for 0 min, 30 min or 120 min. **H** Densitometry analysis of the full-length PINK1 protein bands normalized to the corresponding βIII tubulin bands. One-way ANOVA followed by Tukey’s post hoc test; n = 3; p < 0.0001 (****). Data are expressed as mean±SEM. All data points correspond to ≥3 biological replicates (**H**). Scale bars, 10 µm for whole soma, 2 µm for insets.

**Fig. S4.**
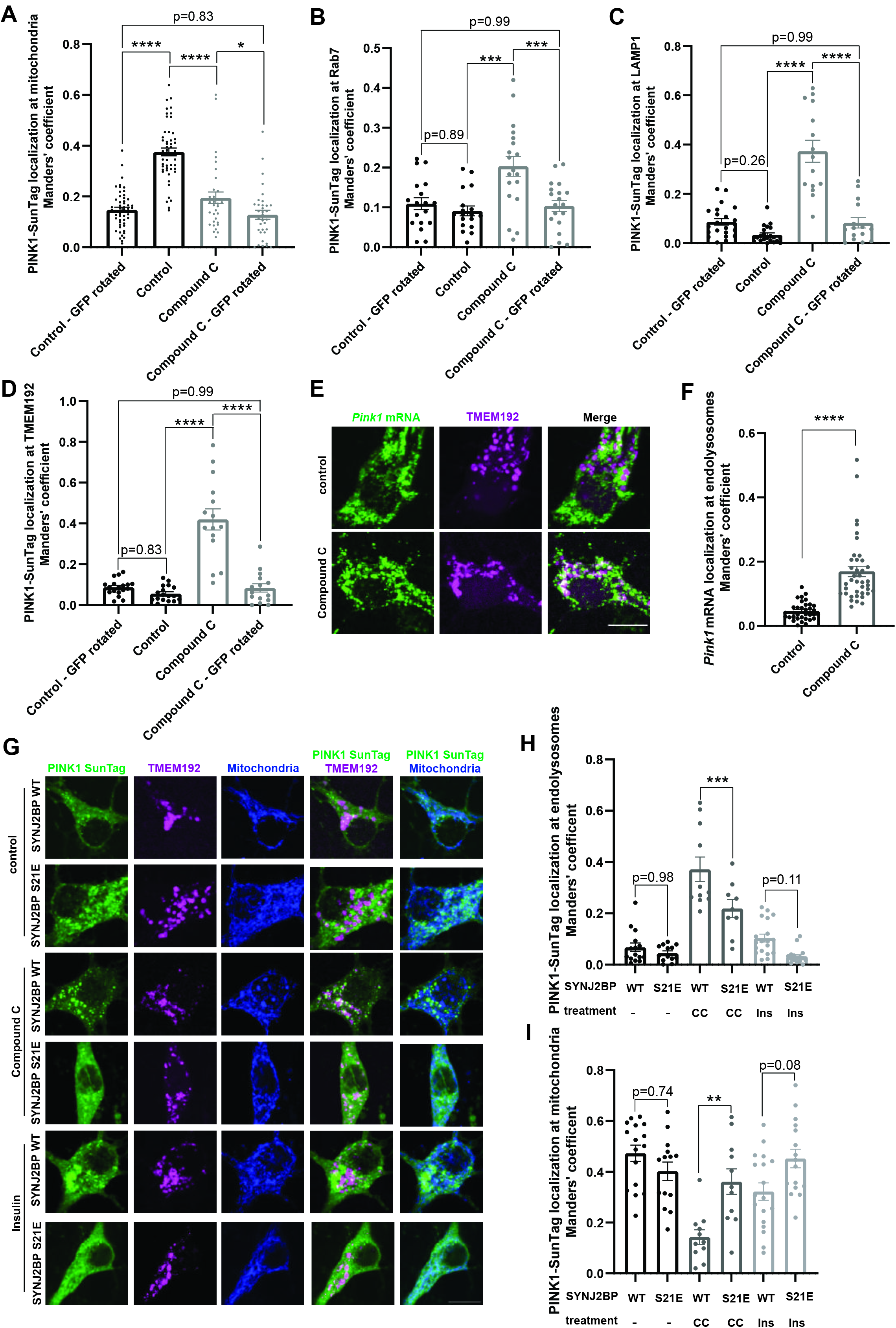
*Pink1* mRNA localizes to endolysosomes upon AMPK inhibition and endolysosomal localization of PINK1-SunTag clusters is prevented by SYNJ2BP S21E overexpression. **A-D** Quantification of the Manders’ coefficient of the colocalization between the PINK1-SunTag signal and the respective organelle signal in hippocampal neurons upon control (untreated) or CC (20 µM, 2 h) treatment. The analysis was conducted on a 10 by 10 µm square in the soma. ‘GFP rotated’ indicates that the PINK1-SunTag channel had been rotated 90° before quantification. **A** Manders’ coefficient between the PINK1-SunTag and the mito-mRaspberry signal. One-way ANOVA followed by Tukey’s post hoc test; n = 34-53; p < 0.05 (*), p < 0.0001 (****). **B** Manders’ coefficient between PINK1-SunTag and Rab7-mCherry signal. One-way ANOVA followed by Tukey’s post hoc test; n = 18-19; p < 0.001 (***). **C** Manders’ coefficient between the PINK1-SunTag and the LAMP1-mCherry signal. One-way ANOVA followed by Tukey’s post hoc test; n = 15-22; p < 0.0001 (****). **D** Manders’ coefficient between the PINK1-SunTag and the TMEM192-RFP signal. One-way ANOVA followed by Tukey’s post hoc test; n = 15-18; p < 0.0001 (****). **E** Representative images of *Pink1* mRNA visualized by the MS2/PP7-split Venus approach and RFP-labeled TMEM192-positive endolysosomes upon control (untreated) or CC (20 µM, 2 h) treatment in hippocampal neurons. **F** Quantification of the Manders’ coefficient of the colocalization between the *Pink1* mRNA signal and the endolysosomal signal as in **E**. Student’s t-test; n = 34-39; p < 0.0001 (****). **G** Representative images of cytosolic PINK1 protein clusters visualized by the SunTag system, BFP-labeled mitochondria and RFP-labeled TMEM192-positive endolysosomes upon control (untreated), CC (20 µM, 2 h) or insulin (500 nM, 1 h) treatment in hippocampal neurons overexpressing either SYNJ2BP WT or SYNJ2BP S21E. The same set of cells was analyzed in Fig. S2L-M for PINK1-SunTag intensity. **H** Quantification of the Manders’ coefficient of the colocalization between the PINK1 SunTag signal and the endolysosomal signal as in **G**. One-way ANOVA followed by Tukey’s post hoc test; n = 9-19; p < 0.001 (***). **I** Quantification of the Manders’ coefficient of the colocalization between the PINK1-SunTag signal and the mitochondria signal as in **G**. One-way ANOVA followed by Tukey’s post hoc test; n = 11-18; p < 0.01 (**). Data are expressed as mean±SEM. All data points correspond to single cells coming from ≥3 biological replicates (**A,B,C,D,F,H,I**). Scale bar, 10 µm.

**Fig. S5.**
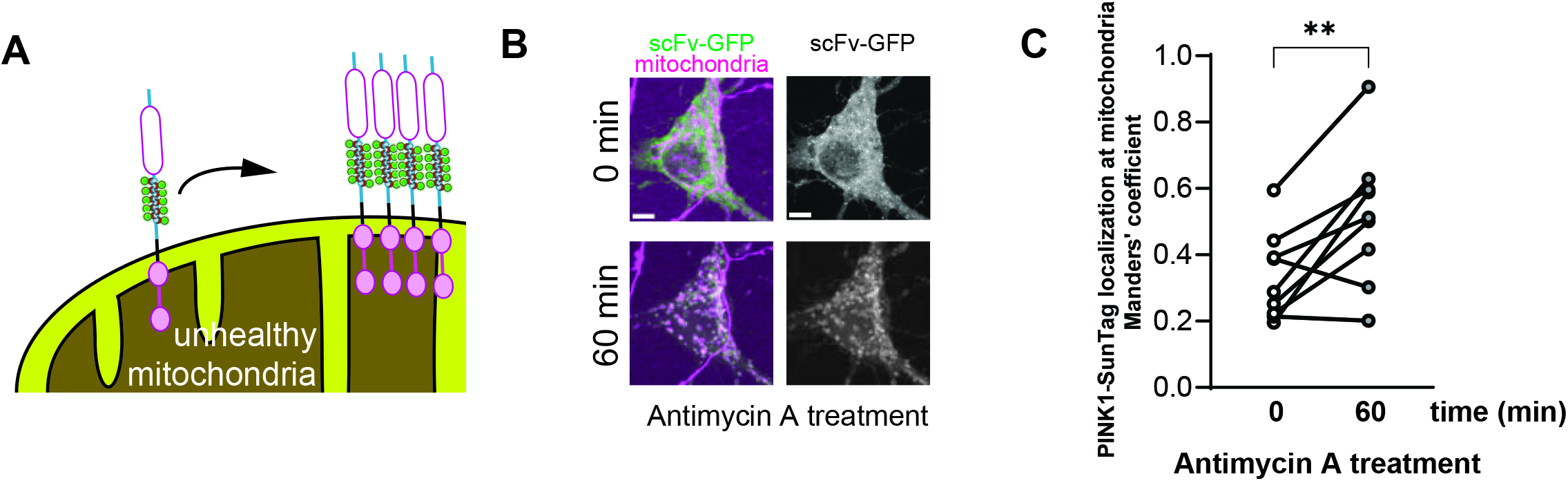
PINK1 SunTag stabilizes around mitochondria upon AA-induced mitochondrial damage **A** Schematic depiction of the accumulation of SunTag nanobodies on stabilized mature PINK1- SunTagcyto upon mitochondrial depolarization. **B** Representative images showing the location of PINK1 protein clusters visualized by the SunTag system (green) and BFP-labeled mitochondria (magenta) before and after incubation with 20 µM Antimycin A for 60 min. **C** Quantification of the Manders’ coefficient of the colocalization between the PINK1-SunTag and the mitochondrial signal as in **B**. Paired student’s t-test; n = 9; p < 0.01 (**). All data points correspond to single cells coming from ≥3 biological replicates (**C**). Scale bar, 5 µm.

**Fig. S6.**
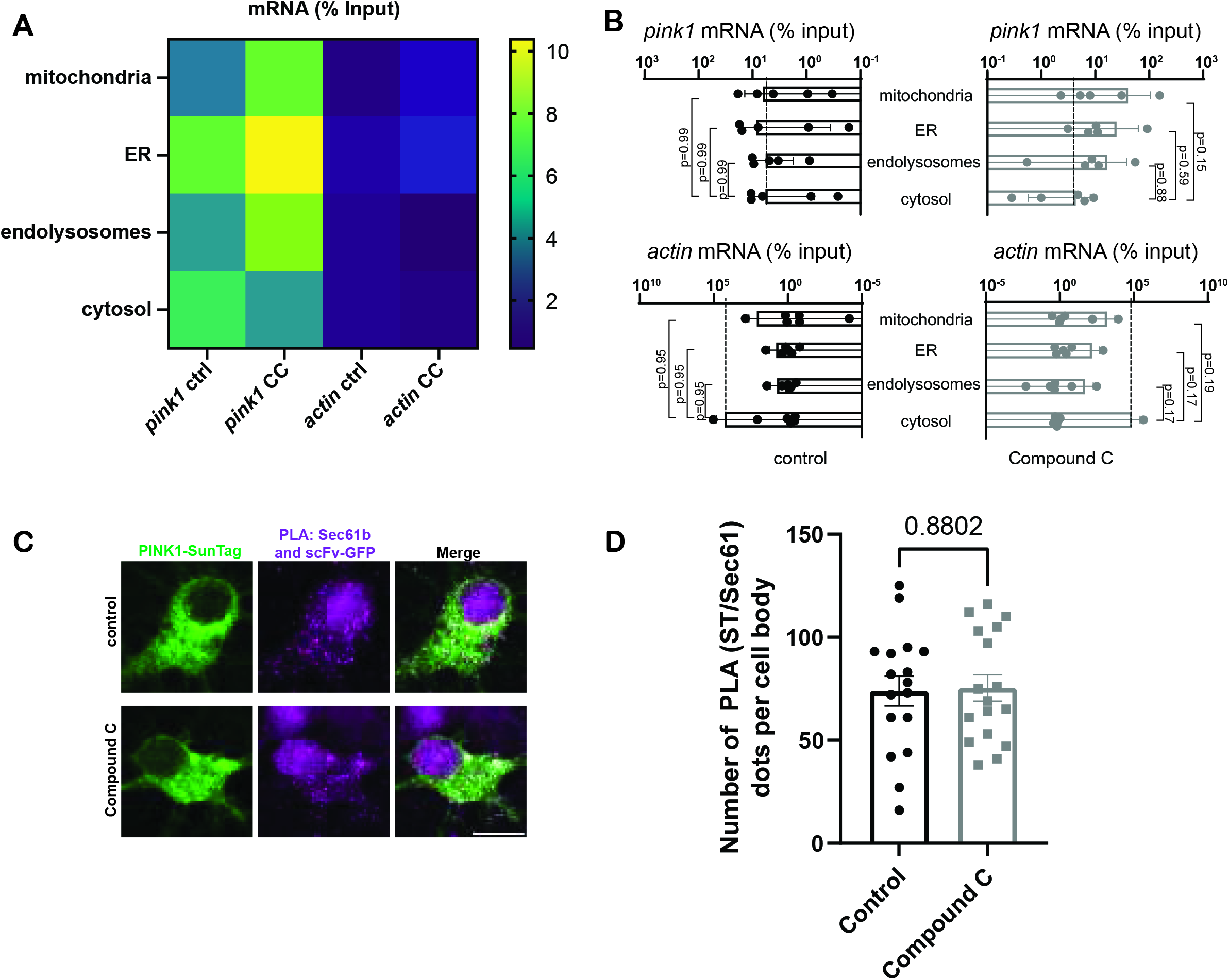
PINK1-SunTag cluster localization to Sec61b does not increase upon CC treatment. **A** Heat-map representing the median value for the enrichment relative to the input of the indicated mRNAs (*Pink1*, *actin*) at BirA-biotinylated ribosomes in the cytosol or in proximity to different organelles (mitochondria, ER, endolysosomes) in mouse cortical neurons, upon control (untreated; ctrl) or CC (20 µM, 2 h) treatment. **B** Enrichment of *Pink1* or *actin* mRNA at BirA-biotinylated ribosomes in the cytosol or in proximity to different organelles (mitochondria, ER, endolysosomes) in mouse cortical neurons, upon control (untreated) or CC (20 µM, 2 h) treatment. Two-way ANOVA followed by Šidák’s multiple comparison test; n = 5-6. **C** Representative images of cytosolic PINK1 clusters visualized by the SunTag system as well as the PLA signal for scFv-GFP and Sec61b upon control and CC (20 µM, 2 h) treatment in hippocampal neurons. **D** Quantification of the number of PLA clusters per soma as in **C**. Student’s t-test; n = 17. Data are expressed as mean±SEM (**B,D**). All data points correspond to either different experiments (**B**) or single cells coming from ≥3 biological replicates (**D**). Scale bar, 10 µm.

**Fig. S7.**
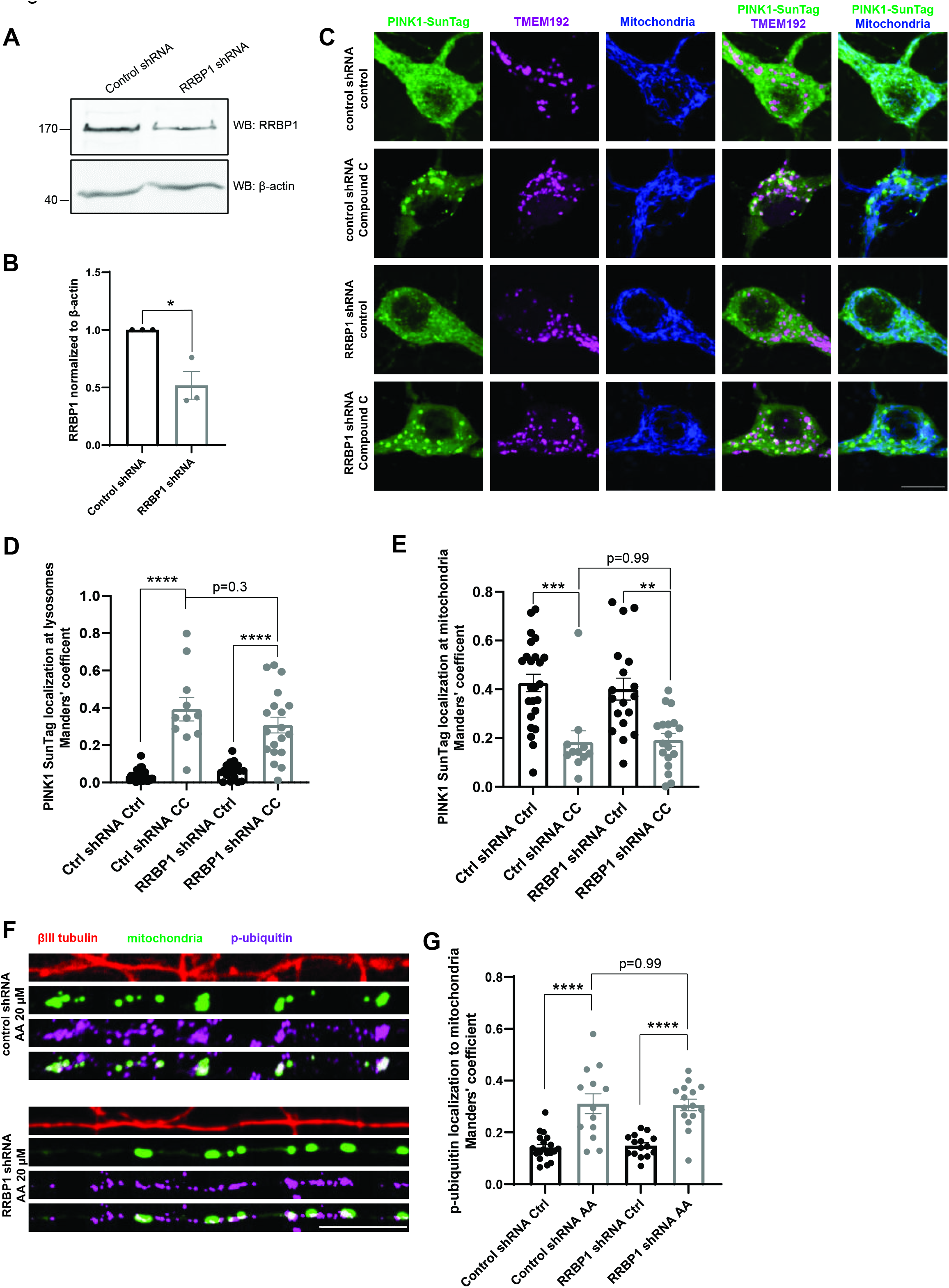
RRBP1 is not involved in the formation of the large PINK1-SunTag clusters upon AMPK inhibition. **A** Representative immunoblot image of lysates from HEK293T cells expressing either control or RRBP1 shRNA. **B** Densitometry analysis of the RRBP1 protein bands normalized to the corresponding β-actin bands as in **A**. Student’s t-test; n = 3; p < 0.05 (*). **C** Representative images of cytosolic PINK1 protein clusters visualized by the SunTag system, BFP-labeled mitochondria and RFP-labeled TMEM192-positive endolysosomes upon control (untreated) or CC (20 µM, 2 h) treatment in hippocampal neurons overexpressing either control (Ctrl) or RRBP1 shRNA. **D** Quantification of the Manders’ coefficient of the colocalization between the PINK1-SunTag signal and the endolysosomal signal as in **C**. One-way ANOVA followed by Tukey’s post hoc test; n = 11-24; p < 0.0001 (****). **E** Quantification of the Manders’ coefficient of the colocalization between the PINK1-SunTag signal and the mitochondria signal as in **C.** One-way ANOVA followed by Tukey’s post hoc test; n = 11-24; p < 0.01 (**), p < 0.001 (***). **F** Representative images of neurites of hippocampal neurons overexpressing mito-meGFP as well as either control shRNA or RRBP1 shRNA. The neurons were treated with vehicle (control) or with Antimycin A (AA, 20 µM, 45 min) and immunostained against phospho-ubiquitin and βIII tubulin. **G** Quantification of the Manders’ coefficient of the colocalization between the phospho-ubiquitin and the mitochondrial signal as in **F**. One-way ANOVA followed by Tukey’s post hoc test; n = 13- 19; p < 0.0001 (****). Data are expressed as mean±SEM. All data points correspond to 3 biological replicates (**B**) or single cells coming from ≥3 biological replicates (**D**,**E,G**). Scale bars, 10 µm.

**Fig. S8.**
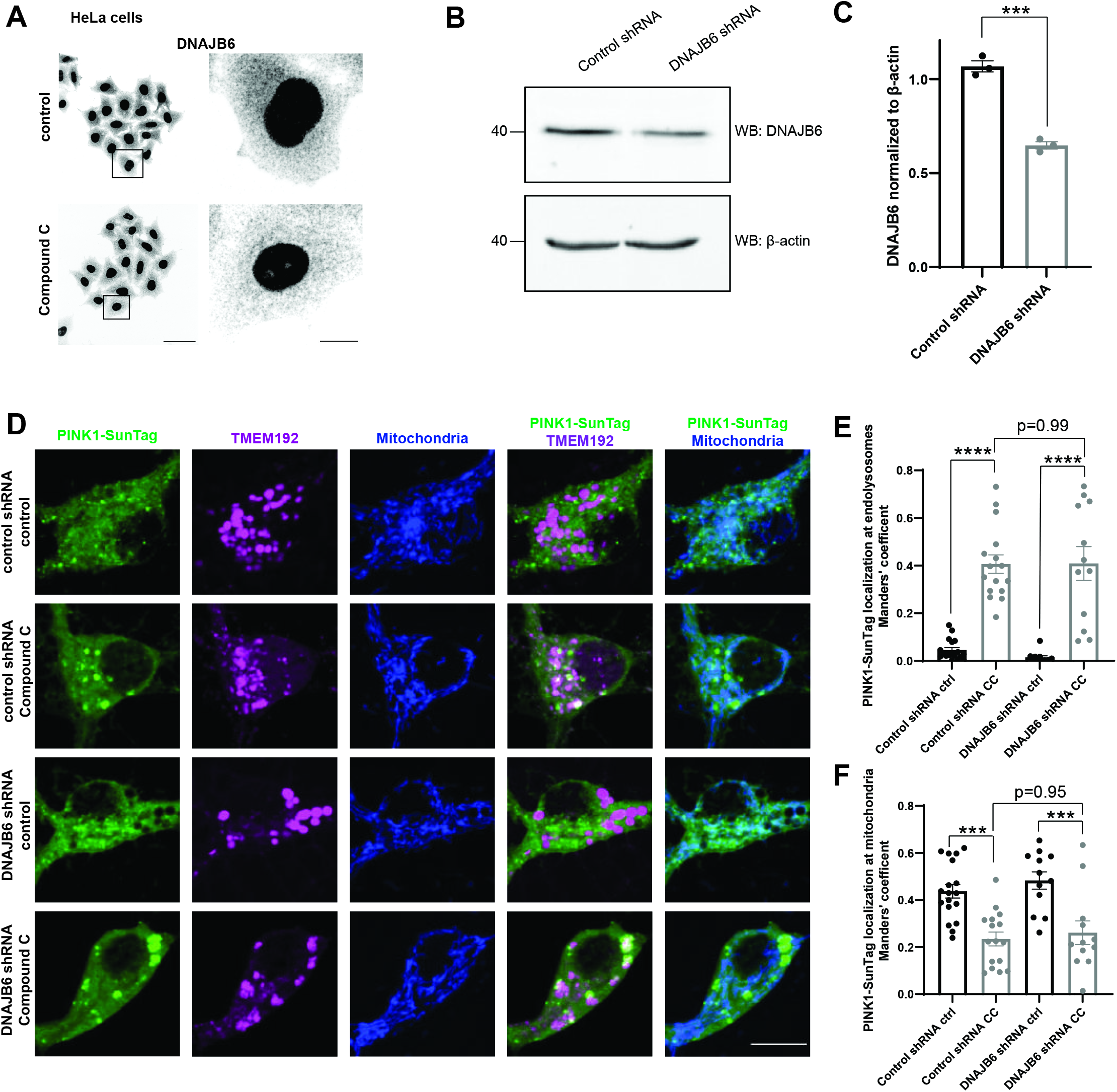
DNAJB6 does not affect localization of cytosolic PINK1-SunTag clusters. **A** Representative images of DNAJB6 immunostaining upon control (untreated) or CC (20 µM, 2 h) in HeLa cells. **B** Representative immunoblot image of lysates from HEK293T cells expressing either control or DNAJB6 shRNA. **C** Densitometry analysis of the DNAJB6 protein bands normalized to the corresponding β-actin bands as in **B**. Student’s t-test; n = 3; p < 0.001 (***). **D** Representative images of cytosolic PINK1 protein clusters visualized by the SunTag system, BFP-labeled mitochondria and RFP-labeled TMEM192-positive endolysosomes upon control (untreated) or CC (20 µM, 2 h) treatment in hippocampal neurons overexpressing either control or DNAJB6 shRNA. **E** Quantification of the Manders’ coefficient of the colocalization between the PINK1-SunTag signal and the endolysosomal signal as in **D**. One-way ANOVA followed by Tukey’s post hoc test; n = 12-18; p < 0.0001 (****). **F** Quantification of the Manders’ coefficient of the colocalization between the PINK1-SunTag signal and the mitochondria signal as in **D**. One-way ANOVA followed by Tukey’s post hoc test; n = 12-18; p < 0.001 (***). Data are expressed as mean±SEM. Data points correspond to biological replicates (**C**) or single cells coming from ≥3 biological replicates (**E,F**). Scale bars, 10 µm for neurons (**D**); 50 µm for HeLa and 10 µm for HeLa inset (**A**).

**Fig. S9.**
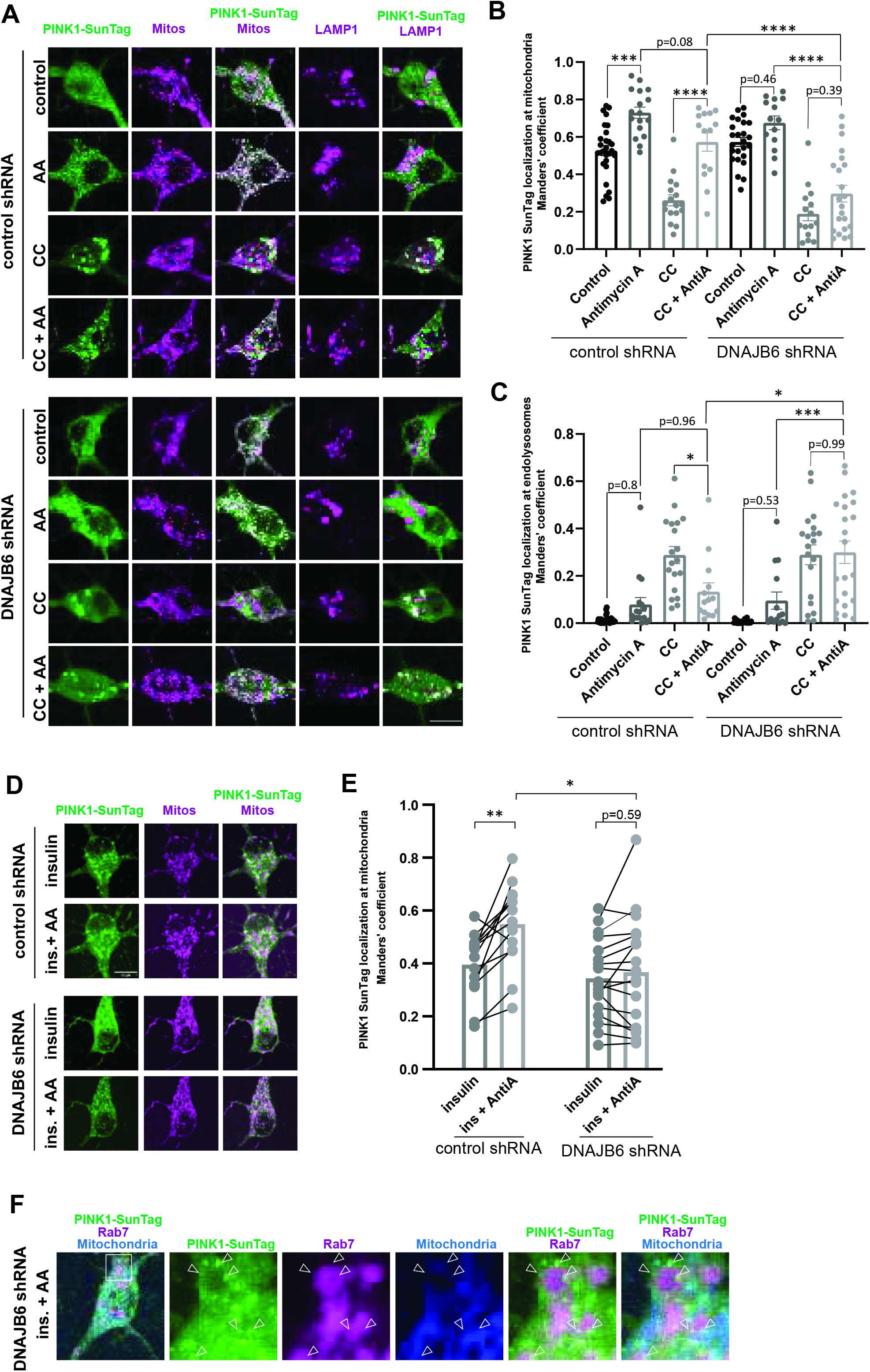
DNAJB6 guides PINK1 from the endolysosomes towards the mitochondria upon induction of mitochondrial damage. **A** Representative images of cytosolic PINK1 protein clusters visualized by the SunTag system and fluorescently-labeled organelles (mito-BFP and LAMP1-mCherry) in the soma of hippocampal neurons expressing control or DNAJB6 shRNA. The neurons were treated with vehicle (control) or AA (20 µM, 45 min), with or without CC pretreatment (20 µM, 1 h). **B** Quantification of the Manders’ coefficient of the colocalization between the PINK1 SunTag signal and the mitochondrial signal as in **A**. One-way ANOVA followed by Tukey’s post hoc test; n = 14-28; p < 0.001 (***), p < 0.0001 (****). **C** Quantification of the Manders’ coefficient of the colocalization between the PINK1 SunTag signal and the endolysosomal signal as in **A**. One-way ANOVA followed by Tukey’s post hoc test; n = 14-28; p < 0.05 (*), p < 0.001 (***). **D** Representative images of cytosolic PINK1 protein clusters visualized by the SunTag system and fluorescently-labeled mitochondria (mito-BFP) in the soma of hippocampal neurons expressing control or DNAJB6 shRNAs. The neurons were treated with insulin (500 nM, 1.5 h) and then co-treated with AA (20 µM, 1 h). **E** Quantification of the Manders’ coefficient of the colocalization between the PINK1 SunTag signal and the mitochondrial signal as in **D**. Two-way ANOVA followed by Tukey’s or Šidák’s post hoc tests; n=14-17; p < 0.05 (*), p < 0.01 (**). **F** Representative images of cytosolic PINK1 protein clusters visualized by the SunTag system and fluorescently-labeled organelles (mito-BFP and Rab7-mCherry) in the soma of hippocampal neurons expressing DNAJB6 shRNA, treated with insulin (500 nM, 1.5 h) and then co-treated with AA (20 µM, 40 min). Arrowheads indicate clusters of PINK1 localized at the endolysosomes. Data are expressed as mean±SEM (**B**) or mean (**E**). All data points correspond to single cells coming from ≥3 biological replicates (**B,C,E**). Scale bars, 10 µm.

**Fig. S10.**
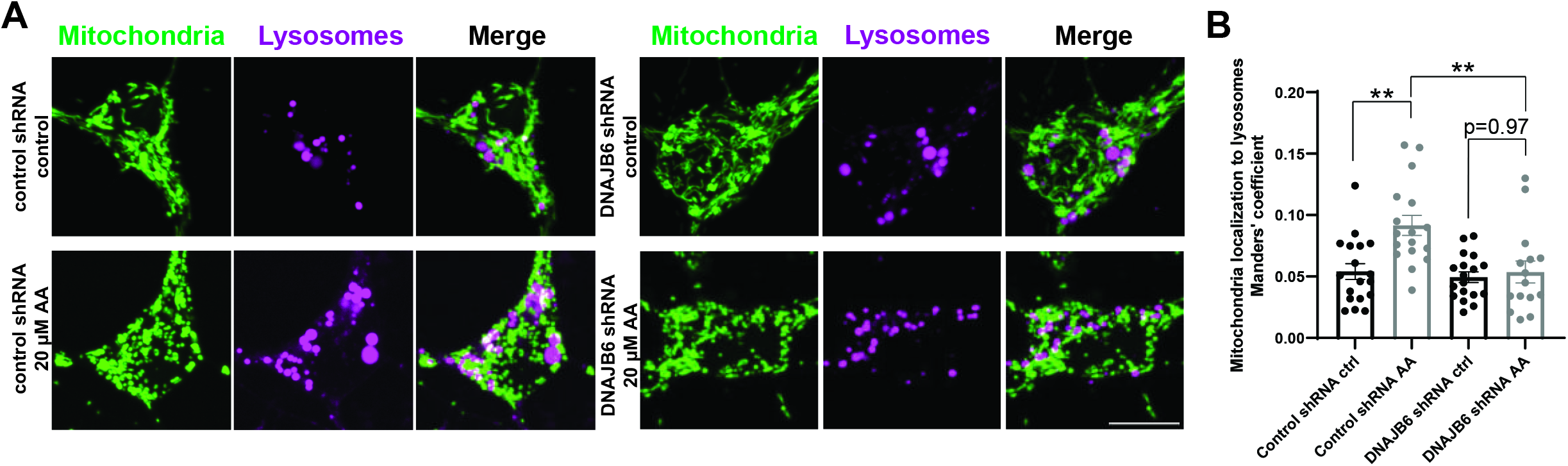
DNAJB6 knockdown impairs localization of damaged mitochondria to lysosomes. **A** Representative images of hippocampal neurons overexpressing meGFP-labeled mitochondria and mCherry-labeled LAMP1-positive lysosomes as well as either control or DNAJB6 shRNA. Neurons were treated with vehicle (control) or with 20 µM AA for 45 min. **B** Quantification of the Manders’ coefficient of the colocalization between the mitochondrial and endolysosomal signal as in **A**. One-way ANOVA followed by Tukey’s post hoc test; n = 15-18; p < 0.01 (**). Data are expressed as mean±SEM. All data points correspond to single cells coming from ≥3 biological replicates (**B**). Scale bar, 10 µm.

## Notes

### Competing Interest Statement

The authors have declared no competing interest.

